# Functional dissection of complex and molecular trait variants at single nucleotide resolution

**DOI:** 10.1101/2024.05.05.592437

**Authors:** Layla Siraj, Rodrigo I. Castro, Hannah Dewey, Susan Kales, Thanh Thanh L. Nguyen, Masahiro Kanai, Daniel Berenzy, Kousuke Mouri, Qingbo S. Wang, Zachary R. McCaw, Sager J. Gosai, François Aguet, Ran Cui, Christopher M. Vockley, Caleb A. Lareau, Yukinori Okada, Alexander Gusev, Thouis R. Jones, Eric S. Lander, Pardis C. Sabeti, Hilary K. Finucane, Steven K. Reilly, Jacob C. Ulirsch, Ryan Tewhey

**Affiliations:** Broad Institute of Harvard and MIT, Cambridge, MA, USA; Program in Biophysics, Harvard Graduate School of Arts and Sciences, Boston, MA, USA; Harvard-Massachusetts Institute of Technology MD/PhD Program, Harvard Medical School, Boston, MA, USA; Analytic and Translational Genetics Unit, Massachusetts General Hospital, Boston, MA, USA; The Jackson Laboratory, Bar Harbor, ME, USA; Department of Genetics, Yale School of Medicine, New Haven, CT, USA; Program in Medical and Population Genetics, Broad Institute of Harvard and MIT, Cambridge, MA, USA; Stanley Center for Psychiatric Research, Broad Institute of Harvard and MIT, Cambridge, MA USA.; Center for Computational and Integrative Biology, Massachusetts General Hospital, Boston, MA, USA.; Department of Statistical Genetics, Osaka University Graduate School of Medicine, Suita, Japan; Department of Genome Informatics, Graduate School of Medicine, the University of Tokyo, Tokyo, Japan; Insitro, South San Francisco, California, USA; Program in Biological and Biomedical Sciences, Harvard Medical School, Boston, MA, USA.; Department of Organismic and Evolutionary Biology, Harvard University, Cambridge, MA, USA.; Howard Hughes Medical Institute, Chevy Chase, MD, USA; The Novo Nordisk Foundation Center for Genomic Mechanisms of Disease, Broad Institute of MIT and Harvard, Cambridge, MA, USA; Program in Computational and Systems Biology, Memorial Sloan Kettering Cancer Center, New York, NY, USA; Laboratory for Systems Genetics, RIKEN Center for Integrative Medical Sciences, Kanagawa, Japan; Harvard Medical School and Dana-Farber Cancer Institute, Boston, MA, USA; Department of Biology, MIT, Cambridge, MA, USA; Department of Systems Biology, Harvard Medical School, Boston, MA, USA; Wu Tsai Institute, Yale University, New Haven, CT, USA; Illumina Artificial Intelligence Laboratory, Illumina, San Diego, CA, USA; Graduate School of Biomedical Sciences and Engineering, University of Maine, Orono, ME, USA; Graduate School of Biomedical Sciences, Tufts University School of Medicine, Boston, MA, USA

## Abstract

Identifying the causal variants and mechanisms that drive complex traits and diseases remains a core problem in human genetics. The majority of these variants have individually weak effects and lie in non-coding gene-regulatory elements where we lack a complete understanding of how single nucleotide alterations modulate transcriptional processes to affect human phenotypes. To address this, we measured the activity of 221,412 trait-associated variants that had been statistically fine-mapped using a Massively Parallel Reporter Assay (MPRA) in 5 diverse cell-types. We show that MPRA is able to discriminate between likely causal variants and controls, identifying 12,025 regulatory variants with high precision. Although the effects of these variants largely agree with orthogonal measures of function, only 69% can plausibly be explained by the disruption of a known transcription factor (TF) binding motif. We dissect the mechanisms of 136 variants using saturation mutagenesis and assign impacted TFs for 91% of variants without a clear canonical mechanism. Finally, we provide evidence that epistasis is prevalent for variants in close proximity and identify multiple functional variants on the same haplotype at a small, but important, subset of trait-associated loci. Overall, our study provides a systematic functional characterization of likely causal common variants underlying complex and molecular human traits, enabling new insights into the regulatory grammar underlying disease risk.

## Main

Genome-wide association studies (GWASs) have successfully linked tens of thousands of loci to complex human traits and diseases^1,2^, providing a glimpse into the biological underpinnings of these phenotypes and motivating the development of targeted therapeutics^2–7^. Pinpointing the exact causal variant at each locus, a critical step for understanding how individual variants and genes contribute to genetic risk^8–11^, has proven a more elusive task^12,13^. Unlike classical Mendelian disease genetics, the majority of loci associated with complex human traits have individually small effect sizes^14,15^ and do not directly modify the sequences governing the splicing of an mRNA transcript or the amino acid composition of a protein. Instead, most trait-associated variant effects are localized to non-coding *cis*-regulatory elements (CREs), such as promoters or enhancers^16–18^. Furthermore, within individual loci, causal variants segregate with nearby polymorphisms, hindering our ability to identify specific causal alleles from linked alleles^19,20^.

Genetic fine-mapping^21–24^ partially resolves these correlations, known as linkage disequilibrium (LD), and has improved our ability to prioritize the variants driving these associations. At current sample sizes (several hundred thousands), only ∼10-20% of associated loci can be resolved to a single variant^25–27^, and large-scale meta-analyses remain difficult to fine-map accurately due to heterogeneity across cohorts^28^. Thus, most loci are not fully resolved and contain *credible sets* (CSs) of variants, where the probability from the fine-mapping model is distributed across a small set of variants. These CSs are smaller and more experimentally tractable than LD windows^25,26^, but genotypic information alone is not sufficient to fully resolve these loci, let alone understand their regulatory mechanisms.

While large-scale experimental catalogs of CREs^29–31^ and other genomic annotations^32,33^ can aid in causal variant identification^34–37^, most causal variants and their molecular impacts remain largely unknown. Even at loci that can be fine-mapped to a single variant, less than one third of these nucleotide substitutions alter specific units of known regulatory syntax^25,38^, such as disrupting TF binding sites. Direct genome editing of elements containing these variants *in vitro* can uncover impacted sequences and even identify nucleotide-specific effects^39–41^, but these methods remain limited in scale, application, and sensitivity^42,43^. In contrast, high throughput reporter assays can systematically profile allelic effects across an entire element with high sensitivity, at orders of magnitude more loci than endogenous approaches, by recreating aspects of transcriptional regulation at CREs^44–48^.

Here, we test the ability of each allele at 221,412 fine-mapped variants from complex and molecular traits to alter transcriptional output across 5 diverse cellular contexts using massively parallel reporter assays (MPRAs)^49–53^. Using this assay, we identify a high precision set of 12,025 variants in CREs with allelic effects in MPRA, representing 26% of the trait-associated non-coding loci tested. We then extensively evaluate the effects of these variants using experimental and predictive measures of function, nominate molecular mechanisms for fine-mapped regulatory variants, and explore questions of regulatory epistasis and multiple functional variant architectures. Finally, we generate experimental maps of sequence-to-function at nucleotide-resolution for 136 fine-mapped variants in 128 CREs, revealing the complex interplay between common regulatory variants and their sequence contexts.

### High-throughput identification of trait-associated regulatory variants

Previous studies have primarily focused on measuring the regulatory capacity of variants associated with a single complex or molecular trait, typically selecting tens or hundreds of variants in strong LD with each reported sentinel (*i.e.* most strongly associated) variant^44,54–63^. Here, we expand upon these prior works by measuring the allelic effects of variants from 95% CSs, resulting in more efficient capture of the true causal variant(s) than LD windows^64^ (**Fig. 1a**). Variants from selected CSs are associated with 48 complex human traits and diseases from European (UKBB) and East Asian (BBJ) populations, or with changes in expression across 17,969 genes from 49 tissues in a diverse American population (GTEx v8)^25,26,49,50,52^. The complete experiment contains 221,412 fine-mapped variants, including those from 89,387 CSs where over 80% of the total CS probability is tested and 39,868 unique high posterior inclusion probability (PIP > 0.5) variants from the UKBB, BBJ, or GTEx. In addition, we included carefully matched location (n = 32,534), genomic annotation (n = 39,708), and null controls (n = 14,170) (**Fig. 1b**, **Supplementary Fig. 1a, Supplementary Table 1, Methods**).

**Figure 1.**
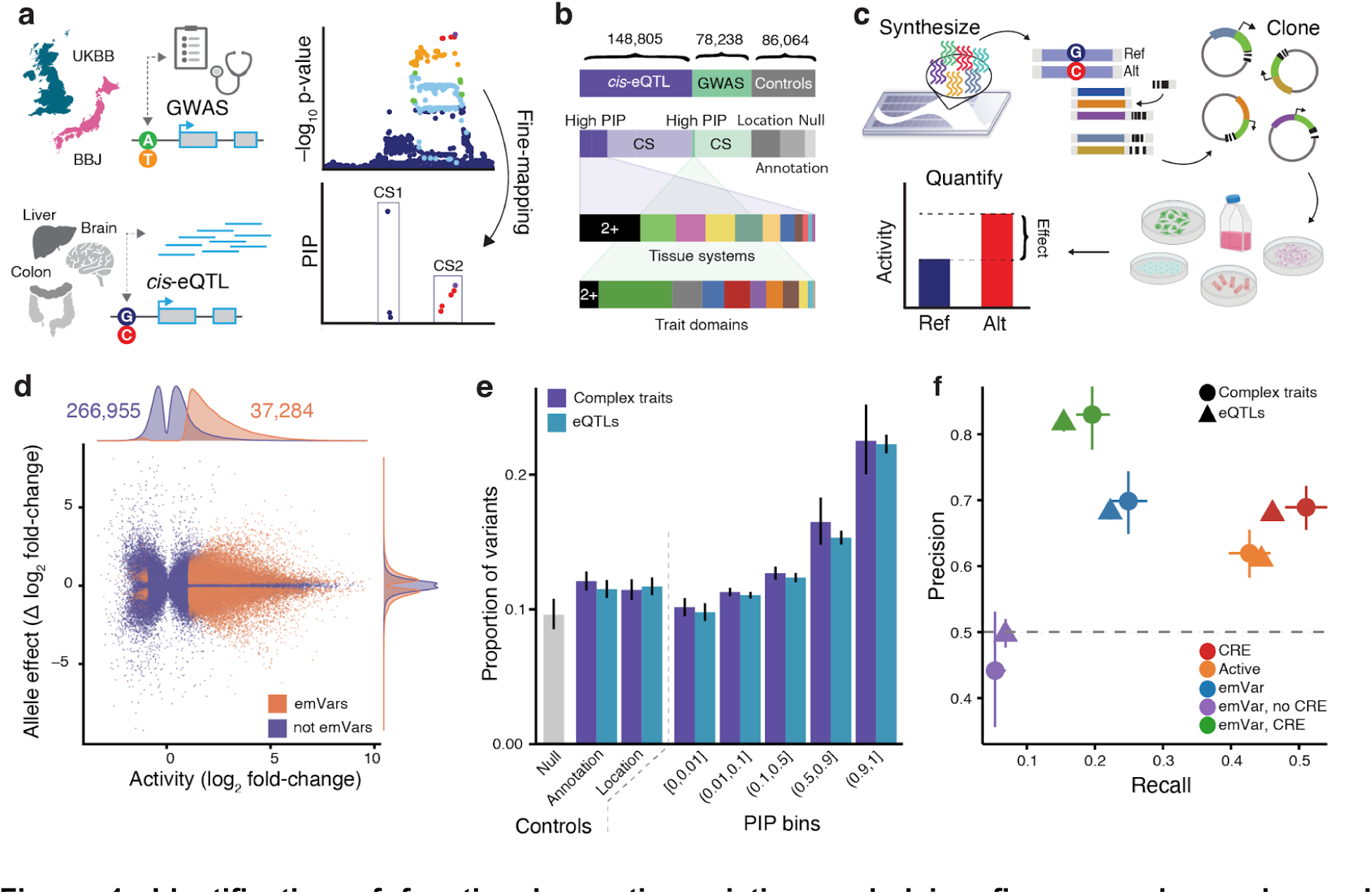
Identification of functional genetic variation underlying fine-mapped complex and molecular traits. **a.** Fine-mapped complex trait variants from UKBB and BBJ, as well as fine-mapped eQTL variants from GTEx v8, were included in this study. **b.** 304,278 unique variants were tested by MPRAs, including 86,064 unique control variants. Nearly 25% of high PIP (> 0.5) eQTLs were associated with gene expression in multiple tissue systems (black), while high PIP complex trait variants were more domain specific. Number of variants per category include overlapping variants, causing totals to exceed the number of unique variants. **c.** Experimental overview of the massively parallel reporter assay (MPRA) experiment. **d.** Element activity and allelic activity results for variants. Each point represents one measurement per variant (selected by best log_2_(fold-change) p-value), with significant expression modulating variants (emVars) denoted in orange, vs non-significant variants in purple. The maximum activity of each variant (ref or alt allele) is shown on the x-axis which has a bimodal activity distribution around 0. Variants with <20 normalized RNA counts are omitted. **e.** The proportion of variants that are emVars across fine-mapping controls and PIP bins stratified by trait type. Error bars represent 95% CIs. **f.** Precision-recall plots evaluating different methods for discriminating between equally-sized sets of positives (PIP > 0.9) and negative (PIP < 0.01) variants. Error bars represent 95% CIs.

A total of 608,556 unique sequences consisting of both alleles across test and control variant sets were designed by centering each allele within 200 bp of its genomic sequence (**Supplementary Table 2**). Sequence oligos were synthesized and cloned upstream of a reporter gene with a minimal promoter and paired with a set of unique 20 bp barcodes in the 3’ UTR (median of 163 barcodes per sequence, **Supplementary Fig. 1b, Methods**). Libraries of MPRA constructs were transfected into four diverse cell-types for each of UKBB/BBJ and GTEx, representing five distinct cell-types overall: blood/myeloid (K562), liver (HepG2), brain (SK-N-SH), and colon (HCT116) or lung epithelial (A549)^44,65^. Barcode sequencing counts expressed from the transfected plasmids were compared to their background representation in the plasmid libraries from at least 5 independent experiments to estimate both the element and allele-specific transcriptional activity of each variant (**Fig. 1c**, **Supplementary Fig. 1c,d, Methods**).

In total, we observed that 92,560 (30.4%) of all tested variants were active (fine-mapped variants: 69,171, 31.2%; control variants: 24,531 28.5%), where at least one of the elements containing an allele impacted transcriptional activity in one or more cell-types (Bonferroni-adjusted P < 0.01 & |log_2_ fold-change [log_2_FC]| > 1), with the majority enhancing rather than repressing transcription (96.6%), reflecting the design of our reporter assay (**Supplementary Table 3**, **Methods**).^44,66^ Of the active variants, 37,284 (40.2%) modulated expression in an allele-specific manner (fine-mapped variants: 28,289, 41.0%; control variants: 9,498, 38.7%, false discovery rate [FDR] < 0.1, **Fig. 1d**, **Supplementary Fig. 1e-h, Supplementary Table 3, Methods**) with predominantly modest effects (median |Δ log_2_ activity| = 0.58). Null controls, location-matched controls, annotation-matched controls, and variants with PIP < 0.01 for all traits had lower levels of element and allele-specific activity compared to causal variants (PIP > 0.9 for at least one trait) which were 2.1-to 2.3-fold more likely to be expression-modulating variants (emVars) than null controls (P = 4.7×10^-17^ and 6.2×10^-50^ for complex traits and eQTLs, respectively, **Fig. 1e**, **Supplementary Fig. 2a**). Allele-specific emVar effects were largely correlated across libraries (Pearson r = 0.82), but background emVar rates varied across libraries and cell-types, reflecting both biological (*e.g.* trait composition of library) and technical (*e.g.* transfection rate) biases (**Supplementary Fig. 2b-c**).

Variants with allele-specific regulatory effects in high-throughput assays have been found to be highly enriched at well-studied regulatory genomic annotations, such as DNase I hypersensitivity, histone modifications, and TF binding sites^44,54,57,62^. However, the sensitivity and specificity of these assays at discriminating between causal and non-causal regulatory variants have typically not been investigated^55–61,63^ or have been estimated in smaller studies with a limited set of “gold standard” variants^44,54,62^. Here, we leverage our recent state-of-the-art fine-mapping studies^25,26^ to select 15,911 unique likely causal variants with PIP > 0.9 for at least one trait and an equal number of matched controls (**Methods**) on which to evaluate our assay. We find that element activity, allele-specific activity, and whether or not the variant lies within a known endogenous CRE (defined as accessible chromatin with activating histone marks, **Methods**) all increase precision compared to chance (0.62, 0.69, and 0.69 respectively) (**Fig. 1f**, **Supplementary Fig. 3a,b, Supplementary Table 4-5**). Importantly, variants in endogenous CREs with allelic MPRA activity have a high precision (eQTLs = 0.82; complex traits = 0.83) and maintain a recall of 0.15 for eQTLs and 0.20 for complex traits, while emVars outside of CREs provide minimal additional information for this task (**Supplementary Table 5**). We note that the exact precision and recall for a specific variant depends upon the underlying trait, assayed cell-types, and resolution of fine-mapping (**Supplementary Fig. 3c-g, Supplementary Table 6**). Overall, we identify 12,025 distinct non-coding, trait-associated variants that significantly alter the regulatory strength of an endogenous CRE sequence in our reporter assay, with 26% of non-coding CSs having ≥1 emVar (**Supplementary Table 7**), including 2,847 with evidence of association in an additional 512 human diseases (**Supplementary Table 8**). In comparison to previous genome-wide approaches^67^, our targeted approach not only reproduced variant effects but also provides improved trait-associated variant coverage, sensitivity, and recall (**Supplementary Fig. 4a-c**).

### Reporter assays recapitulate the native grammar of transcriptional regulation

We have established that coupling MPRAs with measures of endogenous regulatory function can discriminate between causal and non-causal trait-associated regulatory variants with high precision. However, a better understanding of exactly which facets of regulatory grammar are captured by the assay is needed to contextualize any assay-derived mechanisms of non-coding variants underlying complex traits. First, we observe that not only are endogenous distal CREs and promoters more likely to exhibit activity in our assay (odds ratio [OR] = 1.77 and 2.77, respectively, P < 10^-300^), but the magnitude of cell-type agnostic transcriptional activation in our assay correlates well with quantitative measures (maximum) of the chromatin accessibility of DNA across 438 cell-types (**Fig. 2a**, **Methods**).^29^ Of note, the correlation with element activity is higher for variants originating from promoters than for distal CREs (Pearson r = 0.62 vs 0.30). This likely reflects differences in the complexity and composition of TFs binding at distal vs proximal elements^68–71^ as well as aspects of enhancer function in the genome not captured by episomal assays like MPRA^72^. We observe that TFs are more likely to occupy genomic elements containing emVars than those containing high PIP variants alone (mean OR of 3.9 vs 2.1, **Fig. 2b**, **Supplementary Table 9**), with the exception of DMC1^73^, which is involved in recombination, reflecting a type of genome function not measured by our assay. We similarly observe a strong correlation between allele-specific transcriptional activity in our assay and both allele-specific chromatin accessibility and TF occupancy within the genome (Pearson r = 0.65 and 0.54, respectively, **Fig. 2c, Supplementary Fig. 5a**). These findings support the conclusion that our assay is capable of detecting the small, single nucleotide changes that impact transcriptional regulation, and effectively captures the interactions between TF binding and transcriptional activation within the native chromatin environment.

**Figure 2.**
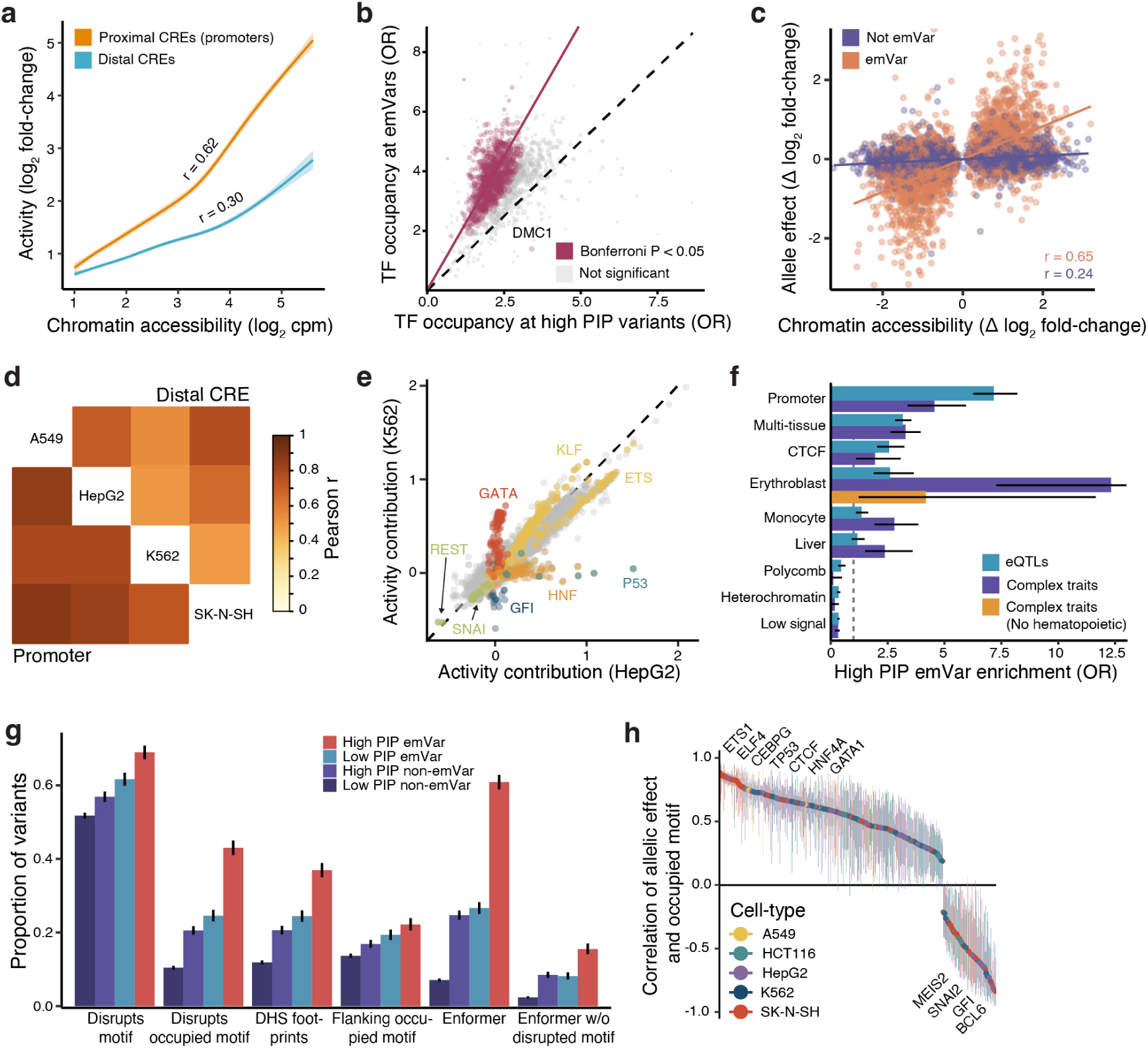
Reporter assays recapitulate endogenous regulatory function. **a.** Correlation (Pearson r) of element MPRA activity with chromatin accessibility at promoters (orange) and distal CREs (blue). Ribbons represent 95% confidence intervals. **b**. TF occupancy at emVars is compared to TF occupancy at high PIP variants for 1,233 TFs. Odds ratios are calculated as emVars vs non-emVars (y-axis) and high PIP (PIP > 0.5) vs low PIP (x-axis). Point size is proportional to the square root of the number of ChIP peaks overlapping variants in this analysis. A linear regression fit through the origin is shown in burgundy. TFs with significant differential enrichment are highlighted in burgundy (Bonferroni-adjusted P < 0.05). **c**. Chromatin accessibility QTL effects sizes are correlated (Pearson r) with MPRA allelic effects. emVars are shown in orange and non-emVars in purple. **d**. Correlation (Pearson r) of element activity at promoters and distal CREs separately across 4 tested cell-types. **e**. Results from a linear regression of normalized motif counts on MPRA activity from 120k CREs in K562 and HepG2 cells. Specific motif families are highlighted in different colors. **f**. Enrichment of high PIP emVars compared to low PIP non-emVars (odds ratio) for complex traits (dark blue) and eQTLs (light blue) in selected genomic annotations defined from SEI.^76^ Error bars represent 95% CIs. **g**. Proportion of variants in each category with the indicated predicted variant effect mechanism. Error bars represent 95% CIs. **h**. Correlation (Spearman ρ) between allelic effects in MPRA and TF binding motif scores for significant TFs (FDR < 0.05). The most significant cell-type is shown for each TF. Error bars represent 95% CIs.

In previous studies, complex trait-associated non-coding variants have been localized most strongly to CREs that are specific to the tissue and cellular programs relevant to their underlying biology^25,65,74^, such as neuronal subsets for Schizophrenia^75^, hepatocytes for circulating lipid levels^25^, and distinct hematopoietic compartments with terminal blood cell production^11^. More recent studies have found important roles for cell-type agnostic elements, such as promoters and multi-tissue enhancers^76^. We investigated whether the allelic effects of non-coding regulatory variants are also primarily cell-type specific when tested by MPRA. First, we observed that the activity of endogenous CREs in our assay is largely correlated across cell-types (median Pearson r = 0.70), is more stable across promoters than across distal CREs (median Pearson r = 0.82 vs 0.61), and is higher at cell-type-matched, cell-type-specific CREs (**Fig. 2d**, **Supplementary Fig. 5b, Methods**). Consistent with these findings, TF binding site contributions to activity are highly correlated between cell-types (r = 0.88-0.94), including both activators (*e.g.* JUN/FOS) and repressors (*e.g.* SNAI, REST). Notably, well-known master regulators, including GATA, KLF, and GFI1B factors in erythrocytes (K562s)^77^, HNF factors in hepatocytes (HepG2s)^78^, and IRF factors in neurons (SK-N-SH)^79^, have cell-type specific effects on activity (**Fig. 2e**, **Methods**). At the allelic level, we find that our assay detects cell-type specific effects, with 25-31% of emVars found in a single cell-type, even after controlling for technical differences in power across cell-types (**Supplementary Table 10**, **Methods**). However, upon closer inspection of annotated regulatory elements^76^, we observe that trait-associated emVars localize not only to cell-type specific^25,65^ but also to cell-type agnostic elements, such as promoters and multi-tissue enhancers (**Fig. 2f**, **Supplementary Fig. 5c, Supplementary Table 11**)^80^. Ultimately, our results provide support for a model where trait associated variants reside in CREs that can be either cell-type specific or cell-type agnostic, the latter being an underappreciated mechanism of causal variants^81^.

### Fine-mapped variants regulate transcription through canonical and non-canonical mechanism

Having established a catalog of likely causal variants acting through gene regulation, we sought to investigate how well existing genomic annotations can nominate the underlying molecular mechanisms for fine-mapped variants. One well-established mechanism is through alteration of the affinity of a transcription factor for a particular binding site by a variant substantially changing the TF binding motif^82^. However, we find that, although 69% of high PIP (> 0.9) CRE emVars disrupt one or more of 839 known TF motifs, 52% of low PIP (< 0.1) CRE non-emVars also disrupt such motifs with seemingly little to no impact on assessed human phenotype (**Fig. 2g**, **Methods**). This is consistent with prior work demonstrating the ubiquity of seemingly non-functional TF motifs across the genome.^83,84^ Instead, we find that at elements with biochemical evidence of TF occupancy, 43% of high PIP emVars disrupt the corresponding known binding motif compared to only 10% of background CRE variants (P < 10^-300^, **Supplementary Table 12**), a 1.03-fold higher enrichment than using overlap with consensus TF footprints protected from cleavage by DNase I. For 685 / 1210 TFs with known binding motifs (57%) and 479 / 614 TF binding motifs with TF occupancy (78%), our assay provides a quantitative readout of the effects of TF binding on gene expression in at least one cell-type (FDR < 0.05, **Fig. 2h**, **Supplementary Table 13**). For example, predicted improvements in PWMs for ETS1 (ρ = 0.86), CEBPG (ρ = 0.83), TP53 (ρ = 0.79), CTCF (ρ = 0.74), and GATA1 (ρ = 0.63) were positively associated with transcriptional activation, whereas changes in SNAI2 (ρ = -0.78), GFI (ρ = -0.83), and BCL6 (ρ = -0.84) were associated with repression. However, in total, we find that no single TF mediates more than 2% of the effects of fine-mapped complex and molecular trait regulatory variants^85^ (**Supplementary Fig. 6a**).

To begin exploring potential mechanisms underlying the remaining 57% of high PIP emVars which do not disrupt an occupied canonical binding motif, we first observe residual enrichment at the flanking regions of occupied TFs (+/-10 bps), suggesting that a subset of trait-associated variants act outside canonical TF binding sites (**Fig. 2g**).^25,38,54^ We thus turned to sequence-based predictions of TF occupancy and chromatin accessibility that can more flexibly model the effects of genetic variation on regulatory element activity.^72,76,86^ Using Enformer^87^, a transformer-based neural network, we created a combined score of variant effects on chromatin accessibility and TF occupancy (**Methods**) and found that this score was the best at distinguishing high PIP emVars from any other category (61% vs 7%, **Fig. 2g**). Of the high PIP emVars not predicted to disrupt a TF motif, the sequence-based model identified 50% as impacting regulatory function (16% of all high PIP emVars, P=7.1×10^-277^ compared to low PIP non-emVars). We investigated which predicted TF occupancy alterations from Enformer best separated high PIP emVars from low PIP non-emVars, finding 66 largely cell-type agnostic factors that were over 4-fold enriched, including RNA Pol II, p300, YY1, ETS1, Jun, and CTCF (**Supplementary Fig. 6b**). Since variant effects from Enformer were often correlated, each high PIP variant is predicted to alter the occupancy of 177 TFs on average, obscuring the exact molecular mechanisms without additional information. Taken together, not only do our results nominate high confidence molecular mechanisms for thousands of fine-mapped regulatory variants, but they provide a lens through which to view the effects of negative selection on trait-associated genetic variation^88,89^, revealing their diverse and often non-canonical regulatory mechanisms^38,54^.

### Proximity between variants increases epistatic effects

After investigating the mechanisms of variants demonstrating allelic effects individually, we next asked how pairs of variants in close proximity may act. The presence of allelic heterogeneity, where multiple independent (different CS) but proximal (same locus) variants are associated with a trait, has been widely established for loci underlying both complex and molecular traits^52,90^. Several studies in other species^91,92^ and recent studies in humans^57,93^ have proposed that multiple causal variants can underlie a single association (single CS), suggesting that there may also be multiple functional variants in tight LD. To directly evaluate such potential genetic architectures in the context of gene regulation, we assayed the regulatory activity for 2,522 pairs of fine-mapped variants residing within endogenous CREs at all 4 possible diplotypes and across 6 different windows (**Fig. 3a**, **Supplementary Fig. 7a-f, Supplementary Tables 14-15, Methods**), including both independent associations (930) and pairs of variants residing in the same CS (1,346). We found that both variants had allele-specific regulatory activity in 46% of pairs, reflecting our prioritization of variant pairs in CREs (**Methods**). Of note, this set contains 210 independent pairs (different CSs) with two functional variants (emVars), suggesting that allelic heterogeneity not only occurs within the same locus but in the same regulatory element.

**Figure 3.**
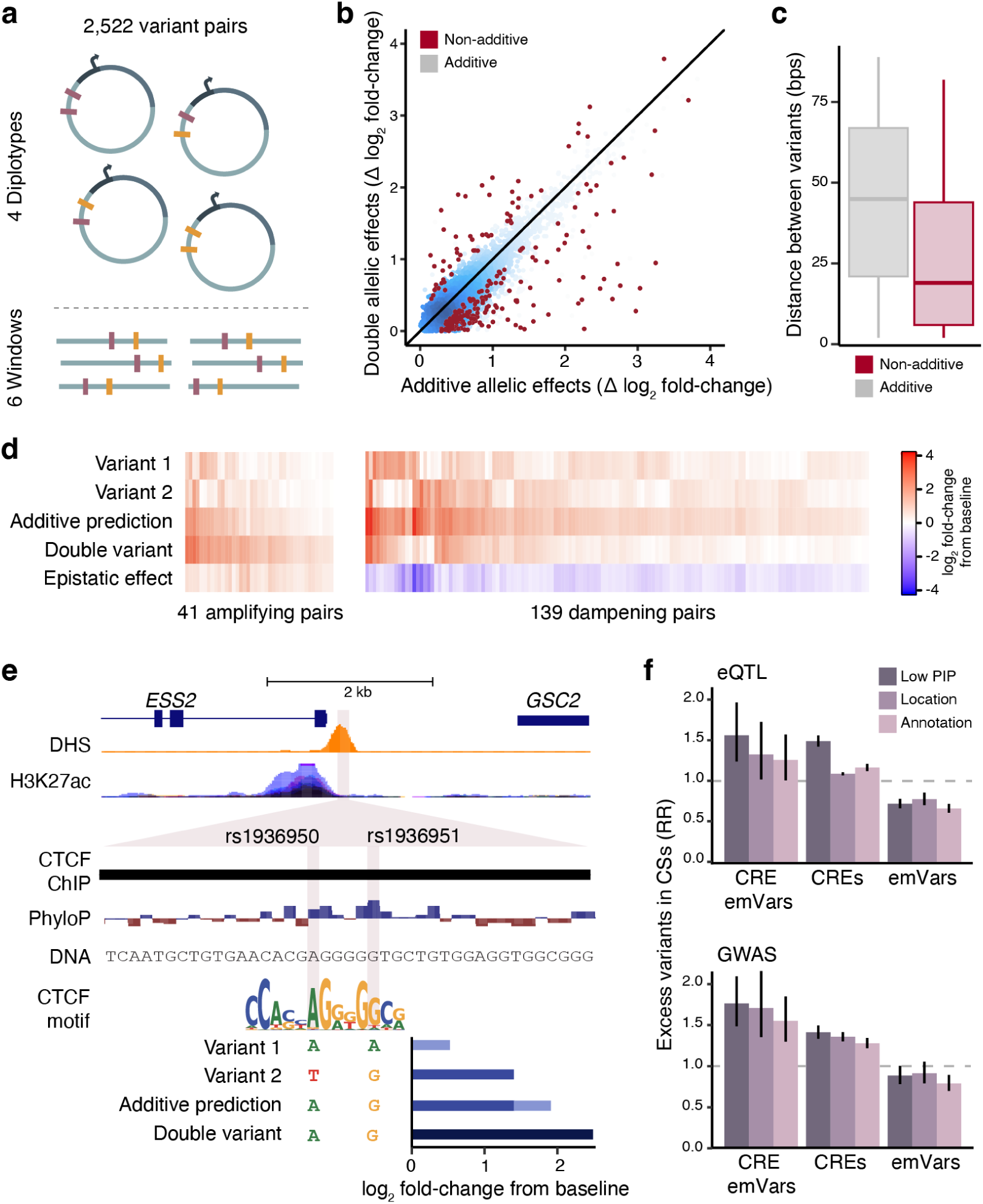
Evidence for regulatory allelic heterogeneity and multiple causal variants. **a**. All 4 diplotypes of fine-mapped variants on the same CRE (< 150 bps, *top*) were tested across six different windows (*bottom*) in MPRA. **b.** Comparison of the expected additive and observed double allele effects after uniformly re-coding diplotypes from largest to smallest effects on activity. Variant pairs with non-additive effects (11%, FDR < 0.05) are shown in red. Each point represents one measurement per variant per cell-type, with the exception of non-additive pairs identified only through a meta-analysis across windows and cell-types, for which one point represents one variant. More dense regions are shown in blue. **c.** Variant pairs with non-additive effects are physically closer than variant pairs with additive effects (p = 10^-8^, Binomial test). Boxes show quartiles, with lines at medians and lower and upper hinges at first and third quartiles; lines extend 1.5 times the interquartile range. **d.** Non-additive pairs of variants are classified into pairs with activating or dampening effects. **e.** Example of an amplifying non-additive variant pair. rs1936950 and rs1936951 (shaded purple) are associated with changes in *ESS2* expression, alter the CTCF binding motif, and fall within a CTCF ChIP-seq peak. Additive prediction is the sum of the re-coded allelic effects from variant 1 (AA vs TA) and variant 2 (TG vs TA). Double variant is the observed difference between (AG vs TA, FDR = 5.5×10^-4^). **f.** Comparison of the number of observed variants in each category (emVars in CREs, CREs, or emVars) and the expected number using three types of controls (location-matched, annotation-matched, or low pip). Risk ratios are from a random effects meta-analysis across experiments (library and cell-type). CSs containing up to 5 variants with r^2^ > 0.9 are included in the analysis. Error bars represent 95% CIs.

GWASs typically investigate only marginal, additive effects, primarily due to their limited power to detect non-additive or epistatic effects^94,95^. Here, we test whether epistatic effects are present at the regulatory level by adding an interaction effect to the emVar model to compare transcriptional activity across diplotypes (**Methods**), finding that 11% of pairs (180) in a CRE with at least one emVar have non-additive effects (FDR < 0.1, **Fig. 3b**, **Supplementary Fig. 7g,h, Supplementary Tables 16-17, Methods**). We observe that epistasis is driven in part by proximity (mean of 26 bp between non-additive variants vs 44 bp for those showing additive effects, P = 10^-10^, **Fig. 3c**) and that most pairs exhibit dampening effects (139/180, Binomial P = 10^-13^), where the element with both activity-increasing alleles has a smaller effect than the sum of individual allele effects (**Fig. 3b,d**, **Supplementary Fig. 7i**). If the relative ratio of dampening to amplifying effects observed here is representative of complex traits as a whole, our results provide evidence for linkage masking^96,97^, where variants escape negative selection by having opposing effects.

We highlight an amplifying epistatic pair example in **Fig. 3e**. rs1936950 and rs1936951 are in near perfect LD, and both are fine-mapped (PIP = 0.64 and 0.36, respectively, and in the same 95% CS) for changes in expression of the *ESS2* gene in the cerebral hemisphere. *ESS2* encodes for DGCR14, in which mutations cause developmental disorders such as DiGeorge Syndrome^98,99^. Each variant’s activating allele significantly increases transcription over baseline (Δ log_2_ activity of 0.51 and 1.41, respectively), but the combined element’s effect is larger than the additive contributions of the alleles (Δ log_2_ activity of 2.54 vs 1.92, FDR = 5.5×10^-4^). The activating alleles of each variant increase sequence similarity to the CTCF PWM, and together they create a near-consensus motif, suggesting that both variants are in fact causal for a single genetic association and, more generally, that TF-driven transcriptional activation is a nonlinear function of TF binding.

Another example of amplifying variants within the same CS is at the *THBS2* locus, a gene with proposed roles in cardiovascular disease and essential hypertension^100,101^. The top two fine-mapped variants, rs9294987 (PIP = 0.48) and rs9294988 (PIP = 0.39), are 2 bps apart and associated with systolic blood pressure in the UKBB. Neither variant is an emVar on its own, with each activating allele only minimally increasing transcription (Δ log_2_ activity of 0.09 and 0.20 respectively). Together, the activating alleles of rs9294987 (C) and rs9294988 (C) substantially increase transcriptional activity of the element (Δ log_2_ activity of 1.68) over the expected additive effect by together creating a near consensus Jun motif **(Supplementary Fig. 7j**). These examples support shared variation potentiation^102^, where effects on a shared regulatory element are not distributed equally but act *on average* to either repress or activate gene expression.

Looking beyond pairs of variants in a single CRE, we continued our investigation into whether single common variant associations (same CS) harbored multiple causal (functional) variants. To rigorously test this hypothesis, we investigated whether we could observe an excess of CRE emVars in CSs containing at least one such variant (**Methods**). Using either low PIP, location-matched, or annotation-matched variants to control for background expectation, we observed significant enrichments for additional CRE emVars within CSs across multiple CS sizes and LD thresholds (risk ratio range: 1.03-2.81, **Fig. 3f**, **Supplementary Table 18**). This corresponds to an excess of CRE emVars in 0.1-3.0% of CSs, which is likely a lower bound when considering the moderate recall of our approach (see **Discussion**). Excesses of CRE emVars were primarily due to variants in multiple enhancers^93^ rather than multiple variants in a single enhancer (range of 0.69-0.83 additional CREs per excess CRE emVar), and were notably higher in CSs that were fine-mapped to smaller sets of variants. Consistent with prior studies, using only CRE annotations^93^, we confirm an excess of fine-mapped variants in these elements (risk ratio range: 1.17-1.60, low PIP controls only) but find that further conditioning on emVar status improves enrichments (**Supplementary Table 18).** Finally, when emVar effects are not restricted to endogenous CREs, we often observe fewer excess emVars than expected (risk ratio range: 0.62-1.39), highlighting the importance of accounting for genomic context and background rate of emVar identification when assessing genetic architectures^44,54,57^.

### High-resolution saturation mutagenesis reveals regulatory mechanisms

We next experimentally interrogated the molecular mechanisms for a subset of trait-associated emVars using MPRA-based saturation mutagenesis (SatMut). We selected 128 CREs containing 136 fine-mapped emVars (PIP range of 0.13 to 1.00) that either (i) disrupt the canonical motifs of occupied TFs or (ii) reside within a CRE but do not disrupt an occupied motif (**Fig. 4a**, **Supplementary Table 19**, **20**, **Methods**). For each allele, we synthesized DNA sequences with every possible single nucleotide mutation (substitution) across the 200 bp sequence and measured their effects on transcription in two cell-types (K562s and HepG2s, **Fig. 4b**). To further investigate haplotype effects, we generated SatMut measurements across all 4 possible diplotypes for 14 pairs of fine-mapped variants, consisting of 22 of the 136 emVars in the test set and 6 additional non-emVars (**Supplementary Fig. 8a, Methods**). All told, allelic effects were estimated for 99% of possible substitutions with an average of 81 unique barcodes and 1,470 reads per replicate. The result is a high quality dataset of >170,000 single nucleotide substitutions across 284 genetic backgrounds that displays excellent concordance with the allelic effects of the original 136 emVars (Pearson r = 0.98, P < 10^-150^, **Supplementary Fig. 8b-d, Supplementary Tables 20, 21, Methods**).

**Figure 4.**
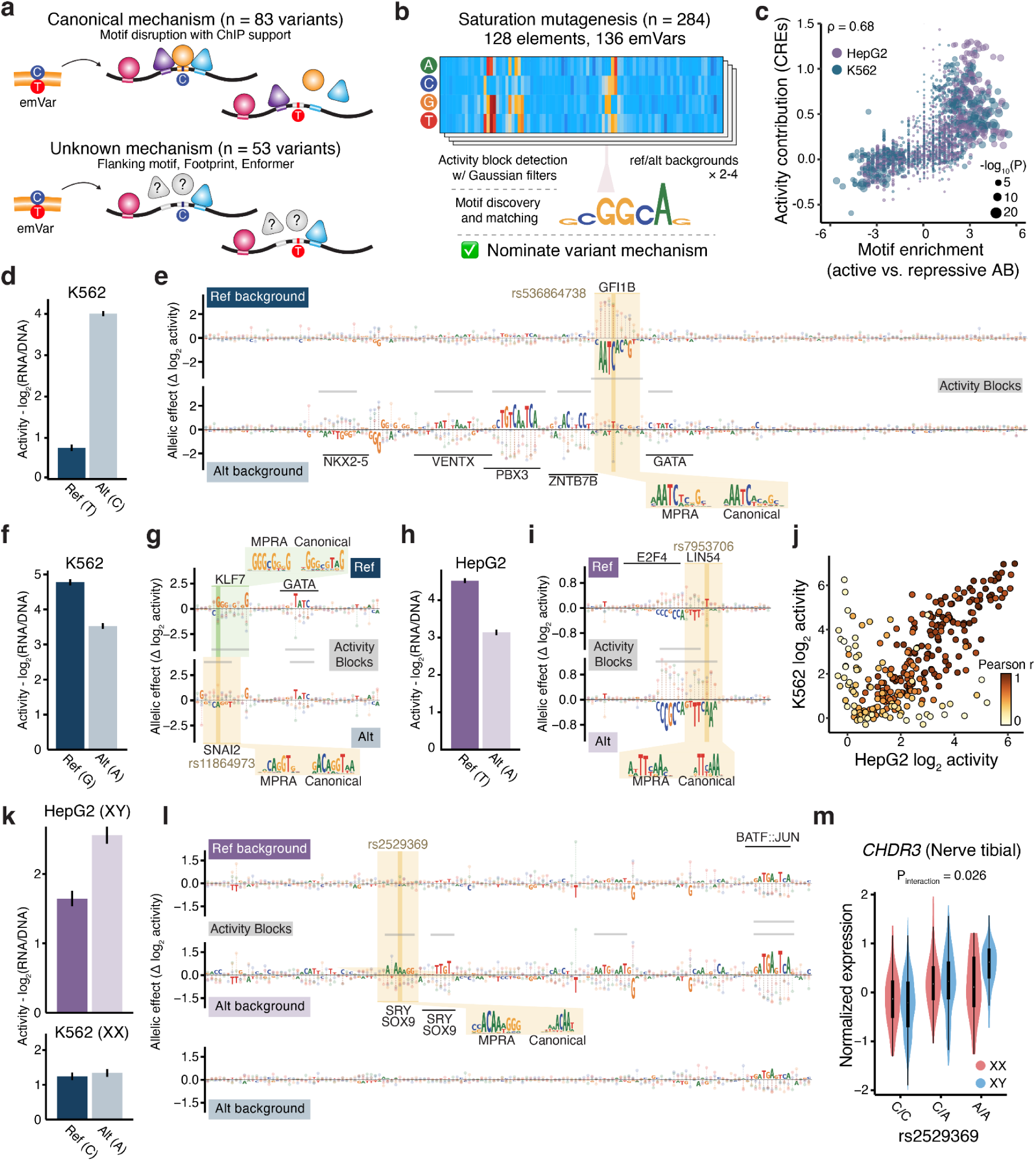
Saturation mutagenesis of 136 fine-mapped emVars. **a.** Schema for two categories included in saturation mutagenesis (SatMut) experiments, emVars with canonical (*left*) or unknown (*right*) mechanisms of action. **b**. Schema of the saturation mutagenesis experiment and analysis. On both allelic backgrounds, mutation of all 200 bases to each of the other three bases is assayed. Short regions that impact transcriptional activity in the SatMut assay are identified as Activity Blocks (ABs) by a Gaussian filtering approach and subsequently matched to motifs PWMs. **c.** Scatter plot of motif effects on activity from CRE sequences presented in Fig. 2e compared to the log_2_ enrichment of motifs in repressive or activating ABs in SatMut sequences by cell-type. The size of each observation corresponds to the p-value from AB enrichment test. **d.,f.,h.,k.** MPRA results of transcriptional activity for the reference (darker shade) and alternative (lighter shade) in either K562 (blues) or HepG2 (purples) for emVars rs536864738 (**d.**), rs11864973 (**f.**), rs7953706 (**h.**), and rs2529369 (**k.**). Error bars indicate SEs. **e.,g.,i.,l.** Nucleotide contribution scores across the 200 bp elements (or zoomed in region) containing emVars from **d., f., i.,** and **l.** are highlighted by a dark yellow bar. Activity measurements for all positions when tested on the reference (*top*) or alternative (*bottom*) allele are depicted as lollipops indicating the change from baseline activity (Δ log_2_ activity). Activity blocks (ABs) are labeled with a gray bar and matching TF motifs are highlighted with a black bar. Shaded boxes overlap allele(s) of interest, with a callout of the SatMut constructed motif (MPRA) and canonical motif PWM (Canonical). **j.** Scatter plot of baseline log_2_ activity for all SatMut tested elements between K562 and HepG2. The correlation between single-nucleotide substitutions for each element is shown. **m**. Violin plot of *CHDR3* expression in tibial nerve tissue from GTEx individuals stratified by both for rs2529369 alleles and sex chromosome status (XX and XY). A significant genotype by sex chromosome interaction is observed (P = 0.026).

Single-nucleotide effects on transcription were physically clustered, suggestive of the disruption of TF binding sites^103^. To identify the short motif-like sequences that affect transcriptional activity based solely upon the results of our saturation mutagenesis assay, we used a Gaussian filtering approach to identify “Activity Blocks” (ABs) of functional nucleotides in each element **(Methods)**. Across all cell-types and background alleles, ABs covered an average of 26% of each element with 98% of sequences containing at least one block. ABs significantly overlapped TF footprints (OR = 1.51, P = 10^-300^), but 44% of ABs and 48% of footprints did not overlap, suggesting shared but non-redundant regulatory features identified by each assay. Since the length of an AB was approximately the size of a TF binding motif (9 bps on average, range of 5-33), we matched position weight matrices (PWMs) for known TFs to PWMs derived from each nucleotide substitution’s contribution to element activity in our SatMut experiments (**Methods**), confidently assigning at least one known TF for 89% of ABs (**Supplementary Table 19**). Notably, this approach more directly implicates both the AB sequence and its assigned TF in regulating transcription from this element than a standard motif search of the DNA sequence alone.

Although each of the 128 selected elements drove transcriptional activity in at least one background and cell-type, a surprising proportion of ABs act as repressors (34%) and exhibited increased expression when disrupted. Known repressor TFs, including GFI1B and SNAI1/3, matched SatMut PWMs exclusively at repressive ABs (**Supplementary Table 21**), and the relative enrichment of TF motifs at activating or repressive ABs was consistent with their inferred direction from the initial screen, validating our initial regression estimates (ρ = 0.68, P < 10^-300^, **Fig. 4c**). Saturation mutagenesis of several emVars disrupting a repressor sequence allowed us to identify activators that are able to drive expression only in the absence of the repressor. For example, the low frequency alternative allele of rs536864738 (MAF = 0.0015) ablates an evolutionary constrained GFI1B repression site when tested in K562 cells, revealing several activators, including a strong adjacent PBX3 site (**Fig. 4d,e**). In the *CCK* promoter, the alleles of the fine-mapped eQTL rs11571842 create either an activating TFDP1 or a repressive ZNF343 sequence, corresponding to increased or decreased *CCK* expression (**Supplementary Fig. 9a,b**). Notably, at the well-known α-globin locus enhancer cluster^104^, our SatMut experiment revealed that the minor allele of rs11864973 converts an activating KLF1 site to a repressive SNAI1 in the MCS-R4 element (**Fig. 4f,g**).

Individual ABs could be attributed to single TFs or combinations of TFs (**Methods**). For instance, SatMut reveals a repressive 16 bp AB downstream of the TSS for *DDX11* whose activity profile most closely matches adjacent E2F4 and Lin54 motifs (**Fig. 4h,i**). Together, these TFs bind DNA with additional cofactors to form the DREAM complex, a potent repressive complex which has been shown to mediate repression of cell cycle genes during G0^105,106^. The reference allele (T) of rs7953706, an eQTL for *DDX11* within this AB, partially ablates this repressive effect, which is supported by recent reports using *in silico* mutagenesis at the region^107^. Lin54 and E2F4, along with 16 other TFs, have evidence of endogenous occupancy from ChIP-seq, but each canonical PWM is either a sub-threshold DNA sequence match (P > 10^-4^) or rs7953706 has only a weak effect on its PWM (Δ PWM < 0.1). In contrast, MPRA SatMut shows that the variant has a measurable impact on transcription by disrupting a repressive element matching adjacent binding sites of TFs making up the DREAM complex.

Between cell-types, single nucleotide substitution effects and AB assignments were often correlated (**Fig. 4j**), with 60% of sequences exhibiting strong correlation (Pearson r > 0.6) or sharing of at least 50% of ABs. Nevertheless, a number of sequences demonstrated clear cell-type specific effects that were largely explained by the presence or absence of GATA motifs for K562 sequences and FOX/HNF motifs for HepG2 sequences. In the previous example (rs536864738), ablation of the GFI1B repressor site only had effects in K562s, but not HepG2s (**Fig. 4d,e, Supplementary Fig. 9c,d**), underscoring the importance of cellular context. A particularly prominent instance of allele- and cell-type specific transcriptional regulation occurs in sequences from a distal CRE (187 kb) for *CDHR3*. SatMut identified two proximal ABs matching SRY/SOX motifs only in the HepG2 cell line and only in sequences with the alternative (A) allele for rs2529369, a fine-mapped eQTL for *CDHR3* (**Fig. 4k,l**). *SRY*, the sex-determining region Y gene, is expressed only from the Y chromosome, which is present in XY HepG2 but not XX K562 cells, and drives expression of *SOX9*, which encodes a TF that has been shown to dimerize and recognize similar endogenous target sites as SRY^108,109^. Since this effect appeared to be dependent on Y chromosome-initiated SRY/SOX9 activity, we investigated whether the sex-dependent allelic effects of rs2529369 could ultimately be observed on *CDHR3* expression in the GTEx v8 cohort. We find that the activating A allele is associated with higher *CDHR3* expression in XY compared to XX individuals (P = 0.026, **Fig. 4m**), supporting the SatMut nominated variant mechanism and the overall ability of MPRA to accurately identify *in vivo* allelic effects.

### Single nucleotide effects and allelic interactions of trait-associated variants

We investigated how well MPRA and other methods could explain the molecular mechanisms underlying fine-mapped emVars. Overlap with an AB or matching motif explained 91% of emVars (124 of 136), regardless of whether the emVar disrupted the motif of an occupied TF (92%, 76 of 83) or lacked a clear canonical mechanism (91%, 48 of 53). Overlap with an AB matching motif explained more emVar mechanisms than any other predictor, including Enformer^87^ (P = 0.001), particularly for non-canonical or unknown mechanisms (**Fig. 5a**, **Supplementary Table 19)**. Notably, SatMut nominated at least one TF for 30 of 31 variants (97%) where traditional PWM matching failed to identify a TF disruption. For most variants (18 of 30) where only SatMut identified a disrupted TF, a predicted PWM disruption could be matched after relaxing the calling thresholds (**Methods**), with the remaining misses partly attributed to the disruption of a low information content motif (*i.e.* short motifs) or the variant residing in a low information content nucleotide.

**Figure 5.**
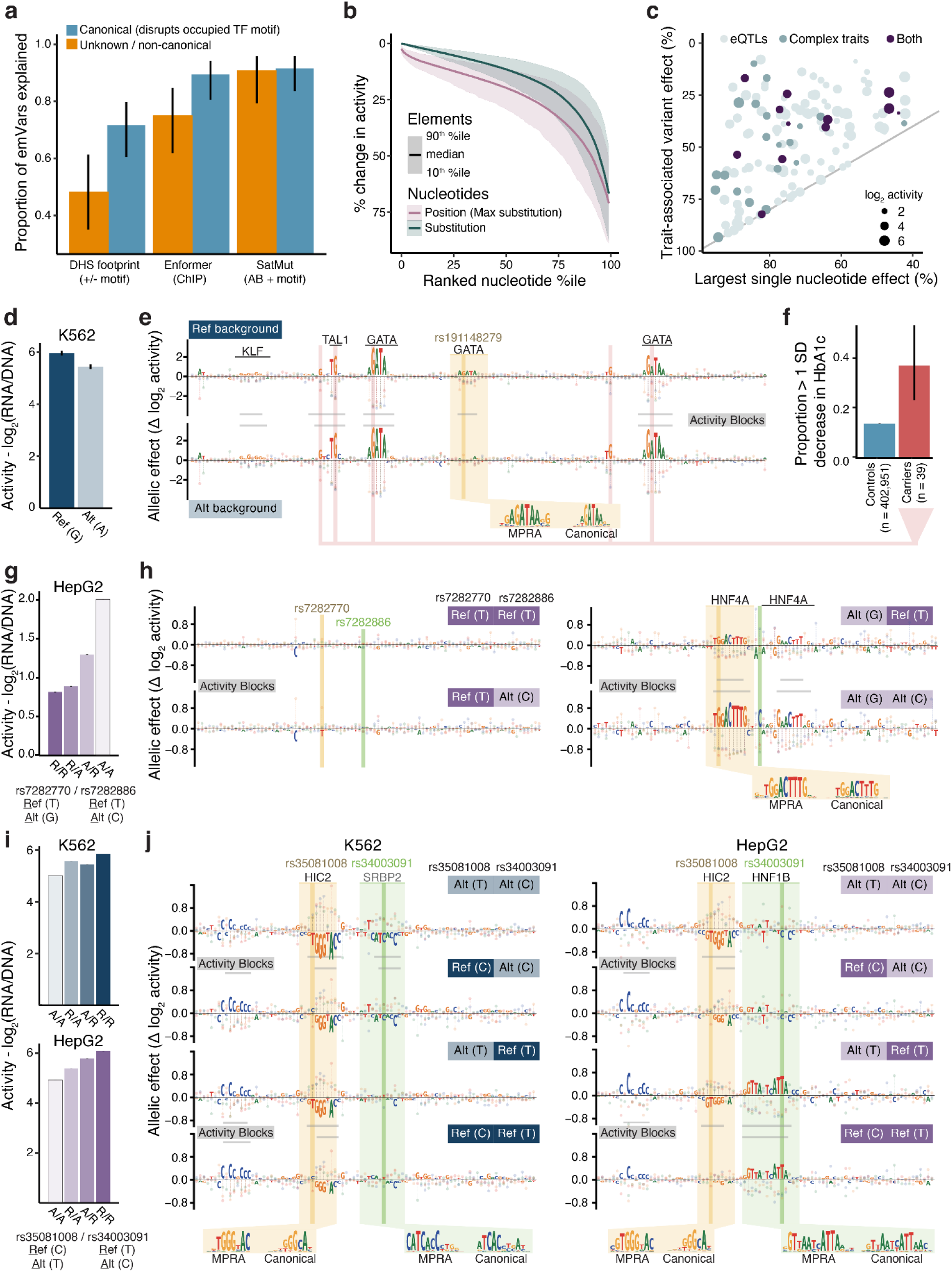
Saturation mutagenesis uncovers canonical, non-canonical, and interacting variant mechanisms. **a**. Proportion of emVars explained by canonical TF mechanisms (blue) or non-canonical mechanisms (orange) across different categories of annotation. Error bars represent 95% CIs. **b**. Cumulative distribution of the percent change in activity for every substitution in each saturation mutagenesis (SatMut) element (teal) or the largest (max) substitution at each nucleotide position (pink). Ribbons represent percent change in activity at the 10^th^ percentile element and the 90^th^ percentile element. **c**. Trait associated variant effects compared to the largest single nucleotide effect seen in a SatMut element. Colors represent the type of trait and size represents the baseline activity of the element. **d**. MPRA results of transcriptional activity for rs191148279, which is an emVar in K562. Error bars indicated SEs. **e**. Nucleotide contribution scores across the 200 bp element containing the fine-mapped variant rs191148279, which are highlighted by a dark yellow bar. Activity measurements for all positions when tested on the reference (*top*) or alternative (*bottom*) allele are depicted as lollipops indicating the change from baseline activity (Δ log_2_ activity). Activity blocks (ABs) are labeled with a gray bar and matching TF motifs are highlighted with a black bar. Shaded boxes overlap allele(s) of interest, with a callout of the SatMut constructed motif (MPRA) and canonical motif PWM (Canonical). Locations of large regulatory effects from rare alleles observed in the UKBB are indicated by pink arrows. **f.** The proportion of carriers of these rare alleles with decreased HbA1C (> 1 SD) are compared to controls. Error bars indicate 95% CIs. **g.,h.** MPRA and SatMut results of transcriptional activity for two adjacent emVars rs7282770 and rs7282886, similar to **d.** and **e.** except for all 4 diplotypes. Error bars indicate SEs. **i.,j.** MPRA and SatMut results of transcriptional activity for two interacting emVars rs35081008 and rs34003091, similar to **f.** and **g.** except for results in both K562 (blues) and HepG2 (purples).

We used our SatMut-based maps of sequence-to-function to explore the spectrum of possible mutation and contrast their regulatory impacts with those of natural, trait-associated variation. Across all altered nucleotides tested by MPRA, a median of 9% of positions and 5% of substitutions had large effects on transcription (> 50% increase or decrease, **Fig. 5b**), most of which affected ABs (72%). Full ablation (log_2_ fold-change < 1) of activity was rare (1% of substitutions) and only achievable in 38% of elements. The majority (80%, 2,277) of large effect substitutions did not have evidence of evolutionary constraint, but mutating nucleotides broadly constrained across mammals^110^ (FDR < 5%) was significantly more likely to have a large transcriptional impact (OR = 7.8, P = 10^-50^). In most elements (63%, 80 / 128), the strength of constraint at each position correlated with the impact of an average substitution on transcriptional activity (FDR < 5%, **Methods**), including a subset (10%, 13 / 128) where MPRA measurements and PhyloP scores were highly concordant (Pearson r > 0.4, **Supplementary Fig. 8f-g, Supplementary Table 22**). Compared to the largest effect substitution, we found that fine-mapped emVars reduced activity less on average (47% reduction compared to 73%), although there was large heterogeneity across elements (**Fig. 5c**), including slightly larger reductions at complex trait-associated elements (P = 0.03).

Elements that harbor larger transcriptional effect alleles than the trait-associated emVar provide an intrinsic opportunity to test whether these alleles, which are likely less common in the population, have similar or more extreme effects on the trait of interest. We observed that 16 out of 136 SatMut elements had at least one substitution that impacted transcription by an absolute difference of 50% more than the fine-mapped variant. One such example is an erythroid CRE that is strong in our reporter assay (98th percentile of active K562 elements) containing rs191148279, which was fine-mapped (PIP = 0.92) for glycosylated hemoglobin A1c (HbA1c) levels^111^ and is predicted to fully abrogate GATA1 binding. We detected an AB overlapping this variant with a SatMut motif matching GATA1 (Pearson r = 0.92), but surprisingly the variant has only a weak effect on transcription (Δ log_2_ activity = -0.49, **Fig. 5d,e**). Instead, GATA motif matches at upstream and downstream ABs contribute substantially more to transcriptional activity (*e.g.* Δ log_2_ activity = -2.93 and -2.42 for similar G>A substitutions, **Fig. 5d**). Each of these stronger GATA1 motifs has AB PWMs matching GATA1 cofactor (TAL1 and KLF1) motifs that are important for stable GATA1 binding and function^112–115^. Collectively, these sites appear to limit the transcriptional effects of mutations at the weaker GATA1 site, a phenomenon known as buffering^116,117^. We searched for variants whose single nucleotide substitutions in SatMut are larger than that of the common variant, rs199148279, in 402,990 unrelated individuals from the UK Biobank who underwent whole genome sequencing. Although there are no other common variants in either of the GATA1 or cofactor sites, we identified 39 individuals who are carriers for one of five rare mutations (allele count < 100) affecting these sites (**Fig. 5e**). Compared to individuals without these mutations, carriers had higher variance (P = 4.7×10^-6^) in HbA1c and were more likely to decrease HbA1c by 1 SD or more (OR = 3.8, P = 6.2×10^-5^, **Fig. 5f**), consistent with lower HbA1c levels observed in individuals with the common allele that affects the weak GATA1 site. Compared to standard PWM-based analysis, SatMut and other mutagenesis approaches expand our capacity to interrogate the impacts of genetic variation across the allelic spectrum within regulatory elements.

Next, we used SatMut to study mechanisms underlying fine-mapped variant pairs in the same CRE with the primary aim of validating and uncovering the sequence determinants of 9 variant pairs with non-additive regulatory effects and 5 additive pairs, each of which is associated with a complex trait. Non-additive effects measured in SatMut and the original experiment largely agreed in terms of effect direction (15/16) and effect size (Spearman ρ = 0.97, P = 9.1×10^-10^, **Supplementary Fig. 8e**), with 5 / 9 validated pairs having large (Δ log_2_ activity > 0.5) amplifying effects and 1 / 9 having a large (Δ log_2_ activity < 0.5) dampening effect. SatMut allowed us to distinguish instances where interactive alleles were within the same binding motif or independent motifs, including validation of the proposed mechanism for the previously described epistatic variant pair, rs1936950 and rs1936951, supporting our initial hypothesis that both variants alter the same CTCF site (**Fig. 3e**, **Supplementary Fig. 9e,f**).

A second example of a non-additive interaction where SatMut provides insight into the mechanism is at a liver-specific eQTL for *DOP1B*, where fine-mapping pinpointed a CS containing 3 potential causal variants in its 3rd intron. Although all 3 variants were in CREs, only rs7282770 (PIP = 0.08) was an emVar (Δ log_2_ activity = 0.38 in HepG2s). SatMut revealed that the minor allele (G) of this variant created two adjacent HNF4 binding sites, separated by 3 bps. Introducing the minor allele (C) of rs7282886 (PIP = 0.08), a CS variant in perfect LD (R^2^ = 1.0) with rs7282770 and located 11 bps away between the two HNF4 sites, resulted in a pronounced non-additive increase (Δ log_2_ activity of 2.25 vs 1.48, **Fig. 5g**). This increase was accompanied by enhanced contributions from both HNF4 sites to transcriptional activity (**Fig. 5h**). Although the 3rd CS variant, rs2246810, which is 3.1 kbps downstream, captured most of the probability (PIP = 0.8), this apparent difference is due to a small reduction in pairwise LD attributable to a single individual’s genotype (R^2^ > 0.99). While we cannot exclude that rs2246810 also contributes to differences in *DOP1B* expression, our SatMut results suggest that rs7282770 and rs7282886 are putative causal alleles for the same eQTL, where both minor alleles facilitate the formation and stabilization of dimeric HNF4 activator sites.

Another example of where SatMut provides insight into the mechanisms of multiple variants in the same CRE is at the promoter of *ZNF329*. Here, the minor alleles of rs35081008 and rs34003091 (R^2^ = 0.99) are associated with decreased expression of *ZNF329* in the liver (PIPs = 0.22) and decreased LDL cholesterol (LDL-C) in the UKBB (PIP = 0.50 and 0.43, respectively). Both variants are emVars and decrease transcription in MPRA (Δ log_2_ activity = -0.30 and -0.82, respectively, **Fig. 5i**). SatMut confirms these effects, but reveals a different mechanism for each variant. The minor allele (T) of rs35081008 creates a HIC2 repressor site in both K562 and HepG2 cells, while the minor allele (C) of rs34003091 disrupts a motif for HNF1, a strong activator in hepatocytes, specifically in HepG2s (**Fig. 5j**). The role of rs35081008 as a causal variant is supported by a recent base-editing study^118^, which found increased *ZNF329* expression after converting the minor allele to the major. However, the likely additional causal role of rs34003091, which has a larger transcriptional effect than rs35081008, and the regulatory mechanisms influenced by both variants are distinctly uncovered by MPRA and SatMut.

Finally, since mutagenesis was performed across entire elements on each allelic background, we investigated the interaction between each of the 114 emVars that were centered on the oligo and all 597 possible substitutions at other positions. We found non-additive effects with 10% of substitutions (|Δ activity| > 25%), of which 58% were dampening effects (**Supplementary Fig. 9g**). Most non-additive effects occurred within ABs or matched motifs (78%), and nearly 1 out of 5 substitutions within an AB had non-additive effects (18% vs 4%, P < 10^-300^). Notably, non-additive effects exhibited a strong positional dependence to the central emVar (35% lower odds per log(bp), P < 10^-300^), consistent with our observations from fine-mapped common variant pairs. Amplifying effects were both 85% more likely to be observed in an AB (P = 2.4×10^-119^) and had stronger positional dependence than dampening effects (P = 5.4×10^-33^). Although these results demonstrate that epistatic effects can be frequently observed between random substitutions, the extent to which selective forces exploit or avoid these alleles to shape complex and molecular traits remains an open question.

## Discussion

We combined state-of-the-art statistical fine-mapping^22,25,26^ with comprehensive experimental validation of non-coding, *cis*-regulatory variants to both identify causal trait-associated regulatory variants and dissect their mechanisms. In total, we tested 221,412 fine-mapped variants associated with hundreds of complex human phenotypes and molecular traits, identifying 12,025 variants that alter the transcriptional activity of annotated CREs. Through careful comparisons with 86,064 unique controls, we conclude that MPRA combined with endogenous CRE annotations can confidently identify causal alleles at an important subset of trait associated loci with high precision (0.82-0.83) and modest recall (0.15 for eQTLs and 0.20 for complex traits). The substantial number of fine-mapped loci identified by MPRA also allowed us to assess foundational mechanisms of causal variation. We find that 31% of likely causal (PIP > 0.9) regulatory variants (CRE emVars) are not confidently predicted to alter binding of a canonical TF motif and 39% are not predicted to alter TF occupancy.

To extend our understanding of variant mechanisms, we assessed the fidelity of canonical assignments of regulatory function and dissected those that were enigmatic via saturation mutagenesis of 136 emVars. SatMut experiments uncovered sequence motifs directly associated with transcriptional outcomes and assigned a known TF for 91% of fine-mapped emVars that initially had a non-canonical or unclear mechanism. Our data suggests the inability to resolve the regulatory mechanisms from DNA sequence alone reflects an unresolved trade-off between the sensitivity and specificity of current methods to discriminate between functional and non-functional binding sites of known TFs. We did identify several examples with more cryptic mechanisms, including variants in the spacer between known TF dimers. Our SatMut approach can be applied to other trait-associated regulatory elements to provide a better mechanistic understanding of regulatory architecture at single-nucleotide resolution and can serve as essential validation data for sequence-based models of regulatory function, extending previous works^103^.

Our study further quantifies the genetic architecture of transcriptional regulation, providing clear evidence for widespread allelic heterogeneity^90^ and relatively infrequent but compelling examples of multiple functional variants underlying a single, fine-mapped association. We only detect multiple functional variants within individual CSs for 0.1-3.0% of relevant non-coding associations, after carefully controlling for background effects (1-15% after adjusting for a recall of 15-20%). In general, we find that regulatory variants in the same CRE are capable of dampening each others’ effects, although in rare instances these variants collaborate to amplify their effects on transcription. We note that our findings may not generalize to all variants, traits, or contexts.

Non-coding loci missed by MPRA likely represent a mix of variants with regulatory mechanisms undetectable by MPRA in the cell-types tested, variants with small regulatory effects below detection limits, and variants where fine-mapping is miscalibrated.^119^ We expect that MPRA in additional cell-types and contexts will improve the precision and recall of causal variant identification, especially when coupled with fine-mapping approaches that can incorporate prior functional effect estimates. At the level of the MPRA assay, we estimate that we can improve recall by 2.5-3.4% per additional cell-type on average (**Supplementary Fig. 3c,d, Supplementary Table 5**), although this will eventually yield diminishing returns. As a result, this work suggests modifications to MPRAs^66,120^ and additional high-throughput functional characterization^121,122^ tools will be needed to comprehensively capture variant effects. We caution that to conclusively demonstrate causality for a cellular or organismal phenotype, genetic manipulation in an endogenous genome alongside appropriate phenotypic readouts must be performed^42,123^.

Overall, we demonstrate that large-scale reporter assays provide a tractable experimental system for nominating and dissecting the functions of trait-associated regulatory variants across three large biobanks. Ultimately, application of variant-centric approaches, including not only MPRA but predictive models based on large perturbation datasets, like the one introduced here, will help bridge the critical gap between variant association and functional mechanisms.

## Methods

### Fine-mapping

We obtained fine-mapping summary data, including PIPs for each variant and 95% CSs, for 48 traits in the UKBB and BBJ from previous studies^25,26^. Detailed methods are available from these studies, but we describe the general approach briefly here. The UKBB is a population-based cohort in the United Kingdom, of which 366,194 “white British” individuals were selected for inclusion (https://github.com/Nealelab/UK_Biobank_GWAS). The BBJ is a large, non-European hospital-based biobank from Japan, of which 178,726 individuals of Japanese descent were included in this study. We selected 48 traits across 12 domains (including an “other” category) from the UKBB and BBJ (**Supplementary Table 1**). GWAS was performed on each using a generalized linear mixed model as implemented in SAIGE^124^ (for binary traits) or BOLT-LMM^125,126^ (for quantitative traits) with the following covariates: age, sex, age^2^, age × sex, age^2^ × sex, and the top 20 genetic principal components. Fine-mapping was then performed using SuSiE^22^ with the GWAS summary statistics and in-sample dosage LD in merged 1.5 Mb windows.

Similarly, we obtained fine-mapping results across 49 tissues from the Genotype Tissue Expression Project (GTEx) v8 from previous studies^25,36,53^ and summarize the approach below. The GTEx project is a tissue-based biobank with genotypes (WGS) and gene expression data (RNA-seq) for 838 individuals across 49 tissues (**Supplementary Table 1**), which are grouped into 11 systems. *cis*-eQTL summary statistics, genotype PCs, and other associated data were obtained as input to fine-mapping, which was performed similar to fine-mapping of complex traits from the UKBB, with the major modification that covariates (including genetic PCs) were projected out of the genotypes prior to LD calculation, since GTEx is a more ancestrally heterogeneous cohort.^52^ When collapsing PIPs across multiple tests into a single estimate, the maximum PIP across traits and/or tissues is used. In addition to individual trait CSs, merged CSs across traits or across tissues were obtained as previously described^26^.

### Variant selection & oligo design

We designed 200 bp oligos centered around the reference and alternative alleles of fine-mapped eQTLs from GTEx v8 and fine-mapped complex traits from the UKBB and BBJ cohorts (**Fig. 1a**). Variants were selected for inclusion if they demonstrated a PIP > 0.5 or greater in any tissue or a PIP > 0.1 in any trait or if they fell within a 95% credible set with fewer than 25 variants (eQTLs), 30 variants (quantitative traits), or 75 variants (binary traits). Variants with PIP > 0.1 in any of 3 GTEx tissues approximately corresponding to cell-types for MPRA were also included. That is, we selected additional fine-mapped eQTLs from Liver (matched to HepG2), transverse colon (matched to HCT116), and any Brain tissue (matched to SK-N-SH). All together, we included 148,805 eQTL test variants and 78,238 complex trait test variants (**Fig. 1b**, **Supplementary Tables 1,2**), with 5,645 variants shared across the two sets. All test and control variants (see below) were split into 8 libraries **(Supplementary Table 26)**.

In order to test for potential non-additive effects between trait-associated variants in the same element, we selected 2,522 pairs of variants that were within 150 bps of each other, were fine-mapped with PIP > 0.1 for at least one trait, and overlapped with an endogenous CRE in at least one cell-type (**Supplementary Table 14**). To maximize power to detect non-additive effects, oligos were designed for up to 6 possible windows. Centering on each variant in the pair, the variant was placed at approximately 50 bp, 100 bp, or 150 bp (where the exact location can change for variants causing insertions or deletions), and windows where the second variant was at least 10 bps from the edge of the oligo were included (**Fig. 3a**). For each window, we designed oligos for all four possible diplotypes (Ref/Ref, Ref/Alt, Alt/Ref, Alt/Alt).

In addition to trait-associated test variants, we designed several comprehensive sets of controls. We selected 32,534 “location-matched” controls that were greater than 150 bps but less than 500 bps away from test variants and were not significantly associated with a complex trait (P > 0.0001) or change in gene expression (P > 0.001) in our summary data. Next, we selected 39,708 “annotation-matched” controls that were not in 95% CSs or significantly associated with a trait that were matched for MAF, LD scores, standard genomic annotations (promoter, 3’ UTR, coding, 5’ UTR, intron, CRE, evolutionary constraint) as well as CRE annotations from relevant ENCODE cell-types and tissues. Matching was performed by computing propensity scores and selecting the closest score match using the MatchIt R package. An example of this matching procedure is shown in **Supplementary Fig. 1a**. Finally, we selected 14,710 “null” GWAS controls, where we simulated a typical GWAS phenotype and performed fine-mapping and variant selection in the same manner as was done for complex traits in UKBB and BBJ. To simulate these phenotypes^127^, we drew causal effects from variants present in UKBB or BBJ largely following a previous approach. That is, we assumed 20% trait heritability, either 1,000 (for binary traits) or 5,000 (for quantitative traits) total causal variants, and a MAF-dependent per-allele effect size consistent with previously reported complex trait architecture (α = -0.38)^127^. For all three classes of controls, matching was performed across PIP bins, resulting in 37,437 controls for eQTLs and 48,880 for complex traits (**Fig. 1b**).

We also selected 128 elements containing variants from the original screen that overlap endogenous CREs on which to perform saturation mutagenesis (SatMut). This set contains a total of 136 emVars, where 114 elements each contain 1 emVar, 8 elements contain 2 emVars, and 6 elements contain 1 emVar and 1 non-emVar (**Supplementary Table 19**). Three categories of variants were prioritized for SatMut. First, we prioritized 83 emVars with “canonical regulatory mechanisms”, where a fine-mapped emVar was predicted to disrupt the canonical motif of an occupied TF. Second, we prioritized 53 emVars with “non-canonical or unclear regulatory mechanisms”, where a fine-mapped emVar did not disrupt the canonical motif of a TF with evidence of occupancy from ChIP-seq. Candidate variants were manually inspected and selected variants have at least one alternative measure of function, such as residing within a DHS footprint, lying in the flank (0-10 bps) of an occupied TF motif, or having strong predicted allelic effects in sequence-based models of function, such as Enformer. Third, we prioritized 14 “same CRE variant pairs”, where at least one variant in the pair was an emVar and the pair demonstrated a non-additive effect or both variants were strong candidates for the first two categories. Unless otherwise indicated, the 22 emVars selected as variant pairs were also included in their respective canonical and non-canonical categories.

Overall, variants were chosen in order to survey a variety of complex and molecular traits, a range of PIPs to allow for both confirmation (PIP > 0.9) and exploration (PIP > 0.1 and < 0.9), a variety of predicted effects on different TFs and in different cell-types, and sufficient representation within all 3 categories. In total, we selected 83 variants from category 1 (canonical mechanism), 53 variants from category 2 (non-canonical or unclear regulatory mechanisms), and 14 variant pairs from category 3 (same CRE variant pairs). For each of the 114 single variant elements, we designed 200 bp oligos using both the reference and alternative background sequences, as well as all possible single nucleotide substitutions for both backgrounds (3 alternative nucleotides x 200 positions x 2 backgrounds). For each of the 14 variant pairs, we designed mutagenesis oligos using each diplotype (Ref/Ref, Ref/Alt, Alt/Ref, Alt/Alt) as a background (3 alternative nucleotides x 200 positions x 4 backgrounds).

Finally, we also included technical controls to evaluate the quality of individual MPRA experiments. For each library, we included 91 activity controls, 96 emVar controls, and 506 negative controls selected from previously published experiments^44,128^ (**Supplementary Table 23**). Activity controls were chosen for their ubiquitous activity across multiple cell-types and were selected to represent a broad range of enhancer activities. For the saturation mutagenesis library, we included an additional 2,126 negative controls, selecting 1,876 ORF controls and 250 shuffled sequences from previously published work^129,130^.

### MPRA library construction

The MPRA libraries were constructed as previously described^44^ with several modifications to the original protocol. 230 bp oligonucleotides were synthesized using Agilent Technologies HiFi libraries for complex trait (seven 60K libraries) and eQTL (two 244K libraries) or Twist Biosciences for SatMut (one 300K library) with the designed 200 bp oligo in the middle and 15 bp adaptor sequences on either end. Unique 20 bp barcodes were added by PCR using primers #82 and #202 (**Supplementary Table 24**). Each oligo amplification consisted of 16-32 50 uL reactions containing 25ul Q5 2xMM Ultra II, 2.5uL each of 10uM primer, and 0.5 uL oligo library using the following PCR conditions (98°C for 30s; 6x [98°C for 10s; 60°C for 15s; 65°C for 45s]; 72°C for 5m). PCR products were purified using AMPure XP SPRI (Beckman Coulter, A63881) and incorporated into the SfiI digested pMPRAv3:Δluc:ΔxbaI (Addgene #109035) backbone vector by Gibson assembly (50uL 2xNEB HiFi Assembly MM, 2.2ug DNA oligo pool, 2.0ug SfiI digested pMPRAv3:Δluc:ΔxbaI in 100uL reaction incubated at 50°C for 1 hour). To achieve a library composition of 200-300 barcodes per oligo, a test transformation was performed with 1uL of the 20uL SPRI purified Gibson assembly mixture and 50uL of 10-beta electrocompetent cells (NEB) to determine the optimal amount of Gibson assembly and 10-beta cells to achieve the desired colony-forming unit (CFU) count. Based on the results of the test transformation, a subset of a second identical transformation was taken directly after electroporation and split across ten 1 mL cultures with 10-beta recovery media (NEB) then incubated for 1 hour at 37°C. After 1 hour, each culture was independently expanded in 20 mL of Luria Broth (LB) with 100ug/mL of carbenicillin. In parallel, CFU colony counting plates were created from 4 of the 10 cultures. After 6.5 hours of growth at 37°C, the cultures were individually pelleted and frozen, and the desired number of cultures from 10 expanded cultures was selected, based on the CFU counting plates, to reach an average of 200-300 CFUs (barcodes) per oligonucleotide (15-21 million CFUs for complex traits, 49 million CFUs for eQTLs and 86 million for SatMut.). Cultures were combined and purified using a Qiagen Plasmid Plus Maxi or Midi kit. For each library, sixteen colonies from the CFU plates were checked by colony PCR to determine the oligo insertion rate for the test transformation (insertion rate range: 93%-100%). The expanded purified pMPRAv3:ΔGFP plasmid library was sequenced using Illumina 2 x 150 bp chemistry to acquire oligo-barcode pairings. To construct the final MPRA libraries, 10ug of the pMPRAv3:ΔGFP library was digested with AsiSI, and a GFP amplicon with a minimal TATA promoter (amplified from pMPRAv3:minP-GFP, Addgene #109035) was inserted using Gibson assembly (125uL 2xNEB HiFi Assembly MM, 5.28ug GFP amplicon, 1.6ug pMPRAv3:ΔGFP in a 250uL reaction incubated at 50°C for 1.5 hours). The resulting pMPRAv3 library includes the 200 bp oligonucleotide sequence positioned directly upstream of the minimal promoter and the 20 bp barcode falling in the 3’ UTR of GFP. The Gibson reaction was purified using 1.5x SPRI and eluted in 40 uL water prior to electroporation of 2-16 uL of purified plasmid library into 100-400 uL 10-beta cells. The electroporation was split across six 2 mL cultures and recovered at 37°C for one hour followed by expansion of each 2 mL culture in 500 mL of TB media with 100ug/mL of carbenicillin. After 16 hours of growth at 30°C, plasmid was purified using Qiagen Plasmid Plus Giga Kits. CFU counting plates from 4 of the 500 mL cultures were made to monitor transformation efficiencies (23 million - 185 million CFUs for complex traits, 285 million - 1 billion CFUs for eQTLs, and 123 million - 171 million CFUs for SatMut). Sixteen colonies from the CFU plates were checked by colony PCR to determine the GFP insertion rate for the test transformation with all libraries having a minimum of 80% of plasmid containing a GFP insert.

### MPRA library transfections

We used 5 cell-types for MPRA transfections, selected for representing 5 distinct tissue types. The complex trait libraries were tested in K562, SK-N-SH, HepG2, and A549 cells; the eQTL libraries were tested in K562, SK-N-SH, HepG2, and HCT116 cells; and the SatMut library was tested in K562 and HepG2 cells. All cell lines were acquired from ATCC and routinely tested for mycoplasma and other common contaminants by The Jackson Laboraotry’s Molecular Diagnostic Laboratory. Each replicate experiment was expanded from a low passage cryopreserved aliquot followed by consistent handling for each cell-type across all libraries. Specifically, K562 cells were grown in RPMI (Life Technologies, 61870127) supplemented with 10% FBS (Life Technologies, 26140) maintaining a cell density of 0.5-1 million cells per mL; SK-N-SH cells were grown in DMEM (Life Technologies, 10566-024) supplemented with 10% FBS (Life Technologies, 26140); HepG2 cells were grown in DMEM (Life Technologies, 10566-024) supplemented with 10% FBS (Life Technologies, 26140); A549 cells were grown in Ham’s F-12K (Life Technologies, 21-127-030) supplemented with 10% FBS (Life Technologies, 26140); and HCT116 cells were grown in McCoy’s 5a (Thermo Fisher, 16600108) supplemented with 10% FBS (Life Technologies, 26140). All transfections were performed using a Neon transfection system (Life Technologies), transfecting 150 million (complex traits), 500 million (eQTL) or 700 million (SatMut) cells for each replicate. Transfections were performed using 100 uL tips with optimized settings for each cell-type; K562: 10 million cells per 100 uL, 5 ug of plasmid, 3 pulses of 1450 V for 10 ms; SK-N-SH: 10 million cells per 100 uL, 10 ug of plasmid, 3 pulses of 1200 V for 20 ms; HepG2: 10 million cells per 100 uL, 5 ug of plasmid for complex traits and SatMut or 10 ug of plasmid for eQTLs, 1 pulses of 1200 V for 50 ms; A549: 10 million cells per 100 uL, 5 ug of plasmid, 2 pulses of 1200 V for 30 ms; and HCT116: 10 million cells per 100 uL, 10 ug of plasmid, 2 pulses of 1350 V for 20 ms. We performed a minimum of 5 replicates for each library and harvested each replicate 24 hours post-transfection by rinsing three times with PBS and collecting by centrifugation. Cell pellets were either frozen at -80°C for later processing or homogenized immediately in RLT buffer (RNeasy Maxi kit) and dithiothreitol, prior to freezing at -80°C. For each cell-type and library, no more than two replicates were performed on the same day and with parallel replicates processed from independently expanded batches of cells.

### RNA isolation and MPRA RNA library generation

Total RNA was extracted from the cell homogenates using the Qiagen RNeasy Maxi kit followed by a 1 hour Turbo DNase treatment (Thermo Fisher) with 5 uL of DNase in 1475 uL total volume for 1 hour at 37°C. To stop the DNase digestion, 15 µL 10% SDS and 150 µL of 0.5M EDTA were added, and the sample was incubated at 70°C for 5 minutes. GFP mRNA was then pulled down using a mixture of 3 GFP-specific biotinylated primers at 0.5 nM (#120, #123 and #126, **Supplementary Table 24**) in 0.2x SSC and 33% Formamide (MilliporeSigma 4650-500ML) and incubated for 2.5 hours at 65°C. Biotin probes hybridized with GFP mRNA were then captured by adding 400 uL of pre-washed Sera Mag Beads (Fisher Scientific) eluted in 500 uL of 20X SSX by agitation at room temperature for 15 minutes. The beads were captured on a magnet and washed once with 1X SSX and twice with 0.1X SSC. After removal of the last wash, 50 uL of water was added and the RNA was treated overnight with Turbo DNase at 37°C. The beads were then collected by magnet and the supernatant removed and purified with 2x RNA SPRI. Complementary DNA was created using SuperScript III (Life Technologies) in a 100 uL reaction with a gene specific primer for the GFP transcript (20 uM primer #19, **Supplementary Table 24**) and a modified elongation temperature (47°C for 80 minutes). GFP mRNA abundance was quantified by qPCR to determine the cycle threshold for each replicate using a QuantStudio 5 qPCR instrument (Reaction mix: 5 uL Q5 Ultra II 2x, 0.5 uL 10uM primers #801 and #802, 1.66 uL 1:10,000 Sybrgreen 1, 1 uL cDNA or GFP mRNA in a 10 uL reaction, Cycle conditions: 98°C for 30s; 40x [98°C for 10s; 62°C for 15s; 72°C for 30s], **Supplementary Table 24**). A standard curve with the final MPRA library (10 fg - 1 ng) was run alongside 1 uL of cDNA and 1 uL of GFP mRNA for each replicate. Any samples showing amplification in the mRNA sample were discarded for having plasmid carryover. Replicates within a cell-type and library were diluted to approximately the same concentration based on the qPCR results, and first round PCR (98°C for 30s; 6-13x [98°C for 10s; 62°C for 15s; 72°C for 30s]; 72°C for 2m) with primers #801 and #802 (**Supplementary Table 24**) were used to amplify barcodes associated with GFP cDNA sequences for each replicate. A second round of PCR (98°C for 30s; 6x [98°C for 10s; 68°C for 15s; 72°C for 30s]; 72°C for 2m) was used to add Illumina sequencing adaptors. The resulting MPRA barcode libraries were spiked with 0.01-1% PhiX and sequenced on an Illumina NextSeq 500 or NovaSeq SP.

### MPRA data processing and main analysis

Data from MPRA was analyzed as previously described^44^ using MPRAsuite which includes MPRAmatch (barcode/oligo pairing), MPRAcount (tag assignment and counting) and MPRAmodel (activity and allelic effect estimates) (https://github.com/tewhey-lab/MPRASuite). Briefly, oligo/barcode pairings were identified by merging paired-end reads into single amplicons using Flash^131^. Genomic DNA sequences and barcodes were extracted and the Genomic DNA element was mapped back to the oligo design file using minimap2^132^. Alignments showing greater than 5% error to the design files were discarded, leaving only high quality alignments which were compiled in a lookup table with their matching barcodes. Following sequencing of the RNA and plasmid libraries, 20 bp barcode tags were assigned to oligos from the lookup table and oligo counts were aggregated by addition across all barcodes. Variants with fewer than 30 DNA counts or with exactly 0 RNA counts in oligos for either allele were excluded from all downstream analysis. In the diplotype analyzes a stricter filter was used, and variants with fewer than 100 DNA counts or fewer than 10 RNA counts across all 4 diplotypes were excluded. When variants were assayed in multiple libraries, the library with the highest sum of the square roots of plasmid counts in reference and alternative sequences was selected.

MPRAmodel uses DESeq2 as the framework for normalization and statistical analysis. For each library, counts were normalized across samples using a “summit-shift” approach which centers the distribution of RNA/DNA ratios for each sample at 1. To determine significant differences between DNA plasmid count and RNA count, a negative binomial generalized linear model (GLM) was used within DESeq2 with independent dispersion estimates for each cell-type and library. We estimate element activity in units of log_2_(fold-change) for RNA counts compared to DNA counts by including a contrast in the design matrix of DESeq2 to compare treatment (RNA vs DNA) and use Wald’s test to evaluate the significance of this estimate, correcting for multiple hypothesis testing within each cell-type and library using Bonferroni’s method. We estimate an allelic effect in units of Δ log_2_(fold-change) (or Δ log_2_ activity) for the difference in RNA counts compared to DNA counts at the Alt oligo compared to the Ref oligo by including a contrast in the design matrix of DESeq2 to compare alleles (Alt vs Ref) and treatment (RNA vs DNA) and use Wald’s test to evaluate significance followed by FDR estimation using Benjamini and Hochberg’s method. Elements passing a specific effect size and significance threshold for RNA vs DNA are defined as active, and variants in active elements passing a specific significance threshold are defined as expression modulating variants or emVars. By performing a grid search as described in the following section, we define elements as active if |log_2_[fold-change]| > 1 and Bonferroni-adjusted p-value < 0.01 and variants as emVars if either the reference of alternative allele element is active and FDR < 10% for a non-zero allelic effect. For specific analyses requiring cell-type agnostic estimates, we perform fixed-effect meta-analyses of activity and allelic effects across cell-types and use these estimates.

For saturation mutagenesis, we used the saturation mutagenesis mode of MPRAmatch which applies increased stringency requirements when pairing barcodes to oligo sequences requiring perfect alignment matches during barcode pairing. Since the majority of the SatMut library consisted of sequences with strong MPRA activity by design, we performed a summit-shift normalization procedure where linear scaling factors were determined only from negative controls sequences. Since the primary goal of the SatMut experiment was to compare the effects of substitutions across an element rather than to detect individual emVars, we employed an Empirical Bayes approach to shrink allelic effect sizes towards 0 based upon how accurately they are measured (*i.e.* number of RNA and DNA reads) (**Supplementary Fig. 8c**). First, we obtain an improved estimate of the baseline activity of the background sequence by pooling the log_2_(fold-change) estimates of oligos between the observed 25th and 75th percentiles of substitutions across this element, assuming that approximately 50% of substitutions will have little effect and the mean activity of these substitutions is an unbiased estimate of true baseline activity. Taking the mean and variance of this set as a normal prior, we use DESeq2 estimates of activity (log_2_[fold-change]) and their corresponding SEs to define a normal likelihood, allowing for closed form posterior inference due to conjugacy. We use this prior and likelihood to obtain a posterior distribution for baseline activity of the background sequence. Next, we use the mean and variance of activity estimates between the 5th and 95th percentiles of substitutions to form a less stringent prior with higher variance, better reflecting the distribution of substitutions with true allelic effects. Using a similar Bayesian approach with this prior, we estimate the posterior activity distribution of each substitution. To obtain shrunken estimates of each allelic effect, we subtract the posterior mean activity for each substitution from the posterior mean baseline activity for the background sequence.

### Grid search for activity and allelic effect thresholds

To find the optimal thresholds for activity and for allelic effect, we performed a grid-search across Bonferroni-adjusted p-value or FDR and effect magnitude, aiming to maximize precision and recall in a set with known positives and negatives. For activity, we used overlap with a cis-regulatory element (CRE) **(Supplementary Table 25)** to approximate our positive set and assigning all other oligos to the negative set, reasoning that sequences originating from accessible chromatin marked by H3K27ac should approximate active transcriptional regulatory elements. We selected equal-sized randomly-selected matched sets of variants in CREs and not in CREs (40,000 each for eQTLs, 25,000 each for complex traits) and calculated the precision and recall across a grid of Bonferroni-adjusted p-value thresholds and activity (log_2_[fold-change]) thresholds **(Supplementary Fig. 1e,f)**. Setting |log2[fold-change]| > 1 and Bonferroni-adjusted p-value < 0.01 optimizes the precision-recall tradeoff and is used as the definition of active throughout the manuscript. For allelic effects, we repeated our grid search, using matched sets of high PIP (> 0.9) variants and low PIP variants (< 0.01) as positive and negative sets. First requiring all elements to be active, we calculate precision and recall across a grid of FDR thresholds and allelic effect (Δ log_2_[fold-change]) thresholds **(Supplementary Fig. 1g,h)**. Based on these results, we define emVars as variants residing in active elements with any magnitude of allelic effect and an FDR < 10% for a non-zero allelic effect.

### Non-additive effect estimation and diplotype analyses

Since alternative and reference alleles for trait-associated variants is arbitrarily coded based on the reference genome, rather strictly by effect direction, disease risk, or minor allele, we re-coded the alleles to best aid in interpretation of MPRA effects. Using MPRA estimates of activity for each of the 4 diplotypes, we re-coded alleles in order from lowest to highest activity, with lowest activity as the reference category. We set the reference category as the “decrease-decrease” pair and the highest value as the “increase-increase” pair, with “decrease-increase” and “increase-decrease” pairs in between. The additive allelic effect is defined as the sum of the individual allelic effect (“decrease-increase” + “increase-decrease”). The observed double allele effect is the allelic effect for the “increase-increase” pair compared to the “decrease-decrease” pair. Interaction effects are the difference between the expected additive allelic effect and the double allele effect, which are estimated in the DEseq2 model and re-coded here. Interacting pairs of variants were detected as described below and classified into amplifying or dampening pairs (**Fig. 3b,d, Supplementary Fig. 7i, Supplementary Table 17**).

For each variant pair, we meta-analyzed using a fixed effect meta-analysis for element and allele-specific activity across all available windows for and additionally across all windows and cell-types. Activity and emVar definitions were the same as single-variant definitions, with the exception of additional alleles (diplotypes). Significant interaction effects between pairs of variants were estimated from either a standard fixed effect meta-analysis model (windows and cell-types are independent), similar to activity and emVar effects, or by using a fixed effect model where the covariance between windows is empirically estimated across all pairs based upon the number of shared positions between the windows. Interaction emVars were defined as pairs where at least one diplotype was active and both the FDR was less than 0.05 and the absolute value of the interaction effect was greater than 0.25 for one of the meta-analyses. In the SatMut dataset, interaction effect estimates were similarly obtained by subtraction of the double allele and additive variant posterior activity estimates and interaction emVars were defined as having an interaction effect greater than 0.415 (25% change in activity). Analyses of the distance between pairs of variants was restricted to pairs where variants were tested in all 3 windows (left, middle, and right) since the number of windows a variant pair is tested in is dependent on the distance between the variants (**Fig. 3c)**.

### Multivariate adaptive shrinkage

To account for differences in power when estimating if element activity or allelic effects are shared across cell-types, we used the Multivariate Adaptive Shrinkage in R (mashr) package.^133^ Since MPRA was performed in 4 cell-types for complex trait variants a 4 cell-types for molecular trait variants, we used only the canonical covariance matrices in the mixture model. We selected variants with PIP > 0.1 that are in CREs and fit the mash model separately for complex trait variants and eQTLs separately. Cell-type specific effects were determined using the local false sign rate (lfsr) < 0.05 (**Supplementary Table 10**).

### Coding and non-coding genomic annotations

Genomic annotations were obtained as described previously.^25,26^ Briefly, we ran the Ensembl Variant Effect Predictor (VEP) v85^134^, selecting the most severe consequence for each variant including coding (missense, LoF, and splice site) and UTR annotations. Non-coding variants were annotated as residing within a promoter (using annotations from the S-LDSC baseline model), a CRE, or having evidence of neither. CREs were defined as accessible chromatin in at least one cell-type and evidence of histone modification at H3K27ac in at least one cell-type. We utilized the combined, normalized, and QC’ed chromatin accessibility and histone modification datasets described in^25^, including DNase-seq, ATAC-seq, single-cell ATAC-seq, and H3K27ac measurements from 7 different atlases^29,30,135–139^. This combined dataset consists of over a thousand cell-types: ROADMAP Epigenomics DNase-seq across XXX cell-types and H3K27ac across 98 cell-types^30^; Meuleman *et al.* DNase-seq across 438 cell-types^29^; Domcke *et al.* single cell ATAC-seq across 54 cell-types^135^; Corces *et al.* ATAC-seq data for 18 hematopoietic cell-types^136^; Corces *et al.* single-cell ATAC-seq across 24 brain cell-types^137^; Calderon *et al.* ATAC-seq for 25 immune cell-types^138^; and ChIP-Atlas DNase-seq for 284 cell-types and H3K27ac for 720 cell-types^139^ to identify CREs (**Supplementary Table 25)**.

For analyses comparing the strength of endogenous CRE activity to MPRA activity **(Fig. 2a)**, only CREs from the Meuleman *et al.* dataset^29^ are used and cell-type agnostic CRE activity is estimated as the maximum normalized DNase I hypersensitivity counts across cell-types. In order to identify cell-type specific CREs, we computed the euclidean norm of normalized counts from the Meuleman *et al.* dataset^29^, selected the maximum value across biosamples representing similar cell-types, and selected CREs with a euclidean norm > 0.2. Within CREs from the Meuleman *et al.* dataset^29^, variants were annotated for consensus DHS footprints across 243 cell-types, which were downloaded from^140^ and lifted over to hg19.

### Precision and recall of MPRA

We calculated the precision and recall of different annotations (*e.g.* emVars) for separating likely causal trait-associated variants from background variants. We approximated the true causal (positive) set of variants by using non-coding, trait-associated variants with PIP > 0.9. Background non-coding variant (negative) sets were defined using either location-matched controls, annotation-matched controls, or trait-associated variants with PIP < 0.01. Equally matched sets of positive and negative complex trait variants (995) and eQTL variants (14,999) were obtained. Precision is defined as the number of true positives divided by the number of variants selected as positive by the annotation. Recall is defined as the number of true positives divided by all positives. All three control sets produced similar estimates (**Supplementary Table 4**), and results using low PIP variants as control have the lowest SEs and are used in the main analysis shown in **Fig. 1f**. In addition to evaluating MPRA measurements and genomic annotations, we estimated precision and recall across CS size (**Supplementary Fig. 3e, Supplementary Table 6**) and specific cell-types used in MPRA or for defining CREs (**Supplementary Fig. 3f,g, Supplementary Table 5**). For cell-type analyses, meta points are estimated as averages of cell-type combination and linear regression is used to predict the impact of each additional cell-type on precision and recall.

### Comparison to genome-wide reporter assays

Measurements of allelic effects using the Survey of Regulatory Elements (SuRE) reporter assay for 5.9 million variants in K562 and HepG2 cells were downloaded from^67^. To compare performance between SuRE and MPRA, we again estimated precision and recall using PIP > 0.9 variants as positives and PIP < 0.05 variants as negatives, but now restricting to variants assayed by both approaches.

### Regulatory quantitative trait loci

Chromatin accessibility quantitative trait loci (caQTLs) and allele-specific TF occupancy quantitative trait loci were obtained from ENCODE^29,141^. Variants with an FDR < 0.25 for allele specific differences were included in comparisons to MPRA allelic effects.

### Transcription factor binding motifs and occupancy

Positional motif matches on oligos were identified using a modified version of FIMO^142^ implemented in motif liquidator, for PWMs from the Homo Sapiens Comprehensive Model Collection (HOCOMOCO, H12CORE^143^) and JASPAR^144^ (2022 release). Motifs shorter than 8 bps or with information content (IC) < 12 were excluded. Sequence matches were called when the absolute percentage (scaled by the difference between the best and worst possible sequence matches scores) and the relative percentage (scaled by the difference between the best and random sequence matches scores) of each sequence was greater than 12, and the significance for this match was < 0.0001. Disruption of a motif required a difference of greater than 10% in the absolute percentage of each sequence match to the PWM. Weak calls with disruptive alleles required only an IC score > 8, no relative percentage filter, P < 0.001 and an allelic difference of only 5% in the absolute percentage of each sequence. Motifs were defined as flanking variants if the motif started or ended within 10 bps of the variant **(Fig. 2g).**

TF occupancy for tested elements was determined by ChIP-seq support. We obtained ChIP-seq peaks from ChIP-Atlas, which uniformly reprocessed peaks from 25,823 experiments in 1,046 human cell-types or tissues for 1,741 TFs.^31,139^ For each specific motif match or disruption, a corresponding TF was defined as occupying an element if that element overlapped with a TF peak for any TF that matched a highly similar motif (Pearson r > 0.9 for PWM similarity).

Motif contributions to the activity of elements derived from CREs containing trait-associated variants was estimated using linear regression, modeling the marginal effect of the number of motifs in each element on the activity in K562, HepG2, and SK-N-SH cell types. Prior to fitting the linear regression, activity was quantile normalized across cell-types and a generalized additive model using smoothing splines of 1-mer and 2-mer counts in each element to model activity (each cell-type measurement was an independent observation) was fit using the mgcv R package, and residual activity was used as the outcome for linear regression. P-values for each motif are obtained from a Wald test and FDRs are estimated using Storey’s q-value. Coefficients from these regressions for motifs present in > 0.1% of elements are shown in **Fig. 2e**, although these estimates should not be considered causal effects. We also assessed whether changing the extent of PWM disruption for each motif was correlated with the extent of change in activity in each cell-type. Restricting to variants that are emVars in at least one cell-type and and motifs represented by at least 10 variants, we computed Spearman correlations and estimated FDRs using Storey’s q-value.

### Regulatory sequence classes (Sei)

We obtained Sei sequence classifications and lifted the annotations over to hg19^76^. Variants were excluded if they fell into multiple Sei annotations after liftover (< 1%). The proportion of emVars for each complex trait, stratified by domain, is estimated and shown in **Supplementary Fig. 5c**. Odds ratios were calculated for high PIP (> 0.5) emVars vs low PIP (< 0.1) non-emVars and p-values were obtained from a Fisher’s exact test for both complex trait variants and eQTLs (**Fig. 2f**).

### Enformer

In order to capture more regulatory mechanisms than those mediated through canonical TF motifs, we scored each variant (after lifting over to hg38) using Enformer^87^, a transformer-based neural network model trained to predict functional genomic data, including TF occupancy and chromatin accessibility. Allelic effect predictions across 1,447 distinct tissue / TF pairs and 320 accessible chromatin predictions were z-score normalized to variants with non-emVars in CREs with PIP < 0.05. A combined score for each variant was computed by squaring its allelic effect for individual TF occupancy or accessible chromatin predictions and taking the sum. A threshold for this score was set such that only 5% of non-emVars in CREs with PIP < 0.05 had higher scores (**Fig. 2g**). Thresholds for individual Enformer predictions were similarly obtained.

### Multiple causal variants

We designed a hypothesis test to evaluate whether our data provided evidence of multiple causal variants within independent signals of genetic association (same CS). First, we restricted our analysis to non-coding 95% CSs where we obtained sufficient MPRA measurements on variants harboring most of the CS probability (> 90%). Then, we removed CSs without evidence of the functional annotation being tested (CRE emVars, CRE, or emVars). We counted the total excess number of variants in each remaining CS (containing at least one variant with a functional annotation) and compared this to the background rate estimated from controls (location-matched, annotation-matched, or low PIP), computing both risk ratios and risk differences. Since the emVar call rate has both biological and technical variance across experiments, we designed each library so that individual traits were fully contained within a single library along with all three types of matching controls. Thus, risk ratios and risk differences were computed within each experiment and meta-analyzed using a random effects model implemented in the R package metafor. Meta-analysis was performed separately for complex traits and eQTLs across multiple CS sizes (5, 10, 15) and r^2^ thresholds (0.8, 0.9, 0.99).

### Activity blocks

To identify blocks of quasi-contiguous non-zero variant positions, we used the crossover points between a threshold *T*∈ℝ and a smoothed signal *S* = [*S_k_*∈ℝ] of the Δ log_2_(fold-change) [Δ*_k_*] of an element as follows. For any given sequence element, each Δ*_k_* is obtained by averaging all three posterior allelic substitution effects at each position *k*. Then, *S* is obtained by applying a one-dimensional Gaussian filter (scipy.ndimage.gaussian_filter1d) to [Δ*_k_*] with a kernel standard deviation of *sigma* = 1.15. To identify blocks of negative allelic effects, which indicate positive contributions to activity by the reference sequence, we found the crossover indices {*c_i_*∈ℕ*}* between *S* and *T* = -0.2, and considered as salient those regions [*c_i,_,c_i+1_*] such that

1. *S_k_ ≤ T* = -0.2 ∀*k*∈ [*c_i,_ c_i+1_*], i.e., the signal *S* along the block is below the threshold;
2. *c_i+1_* - *c_i_* +1 ≥ 5, i.e., the length of the region is at least 5;
3. mean(Δ*_k_*, *k*∈[*c_i,_ c_i+1_*]) ≤ -0.15, i.e., the mean of the allelic effects has to be at most 0.15.

Similarly, we took *T* = 0.2 and mean(Δ*_k_*, *k*∈[*c_i,_ c_i+1_*]) ≥ 0.15 to identify blocks of positive allelic effects.

### Motif matches in activity blocks

In order to find a TF motif match to a given activity block *b*=[*b*_start_,*b*_stop_], we constructed a position weight matrix (PWM) from the allelic effects as follows. Let 𝛿*^a^_k_*denote the 3 allelic effects at position *k*∈[*b*_start_,*b*_stop_] between the reference allele *r*∈{A,C,G,T} and the alternative alleles {A,C,G,T}∋*a*≠*r*. We define a matrix *M^t^_k_*, where *t*∈{A,C,G,T} and *k*∈[*b*_start_,*b*_stop_], as *M^t^_k_* = 𝛿*^a^_k_* if *t* = *a*, *M^t^* = 0 if *t* = *r*. Then, we define PWM[*b*, *b*] = 20 * *M^t^* / M for *t*∈{A,C,G,T} and *k*∈[*b*_start_,*b*_stop_], as the position weight matrix induced by the allelic effects in the activity block [*b*_start_,*b*_stop_], where 20 is a chosen temperature parameter and M_max_ = max*_t_*(∑*_k_*|*M^t^*|). Using PWM[*b*_start_,*b*_stop_], we construct the block position probability matrix PPM[*b*_start_,*b*_stop_] by applying the *Softmax* function to the values of PWM[*b*_start_,*b*_stop_] at each position. We also define the block information content matrix ICM[*b*_start_,*b*_stop_] by mapping the values of PPM[*b*_start_,*b*_stop_] at each position into information bits with respect to the uniform background *p_A_*= *p_C_* = *p_G_* = *p_T_* = 0.25. Once we obtained the matrices PWM[*b*_start_,*b*_stop_] and ICM[*b*_start_,*b*_stop_] from the allelic effects in an activity block *b*, we sought to find their best match to a human transcription-factor motif in JASPAR^144^ or HOCOMOCO^143^. Let 𝒟 represent the dataset of known human TF motifs in the union of JASPAR (2022 REDUNDANT) and HOCOMOCO (H12CORE). Let PWM*_m_*, ICM*_m_*, and 𝝀*_m_* represent the position weight matrix, information content matrix, and length, respectively, of a motif *m*∈𝒟. We defined the best alignment offset *o(b*, *m)*∈ℤ between a block *b* = [*b*_start_,*b*_stop_] and a motif *m*∈𝒟 as the integer that maximizes the sum of the element-wise product between ICM[*b*_start_,*b*_stop_] and PWM*_m_* for all possible alignments of the nucleotide positions between the two matrices (with the appropriate zero padding to match the dimensions of the two matrices). Next, we computed the (block) Pearson correlation between ICM[*b*_start_,*b*_stop_] and ICM*_m_* (with the appropriate zero padding) in their best alignment given by *o(b*, *m)* for every *m* in 𝒟, and collected the motifs in the top 50 Pearson correlations. Then, for those top 50 motifs we computed the (match) Pearson correlation between ICM*_m_* and ICM[*b*_start_ + *o(b*, *m)*, *b*_start_ + *o(b*, *m)* + 𝝀*_m_*], that is, the correlation to the ICM allelic effects given the alignment induced by the alignment to the activity block. Finally, we selected the motif with the highest match_Pearson + 0.2 * block_Pearson. If the overlap between the selected TF match and the activity block leaves 5 or more contiguous positions in the block unassigned, we repeat the process above to the unassigned sub-block.

### Rare variant analysis

Whole genome sequencing for 402,990 unrelated (max kinship coefficient < 0.884) individuals in the UK Biobank were analyzed on DNAnexus. Carriers were defined as single nucleotide substitutions with a higher MPRA allelic effect size than rs191148279 from the SatMut experiments in K562 cells. Residual HbA1c phenotypes were obtained as previously described^26^, and an odds ratio using Firth’s correction for > 1 SD change in the phenotype was computed as well as Levene’s test, which was run to test if the variances were different between carriers and non-carriers.

### GTEx interaction analysis

Sex-biased eQTLs were mapped as described in Oliva et al.^145^. Briefly, a linear model with a genotype x biological sex interaction term was used to model normalized gene expression. Genotype principal components and expression PEER factors were included as covariates. The interaction p-value was obtained from a Wald test.

### Constraint

Nucleotide-level PhyloP scores for constraint in 427 mammals were downloaded from Zoonomia^32^. SatMut sequences were lifted over to hg38 and matched to PhyloP scores. Specific positions were defined as constrained if the PhyloP FDR was less than 0.05, corresponding to a PhyloP score > 2.27. Prior to estimating the correlation with PhyloP, Δ log_2_(fold-changes) for all 3 substitutions at each position were averaged and inverted to obtain a contribution score, similar to the height of letters in SatMut examples in **Fig. 4** and **Fig. 5**. Correlation between either the signed contribution score or the |contribution score| and PhyloP using Pearson’s method and corresponding FDRs obtained using Storey’s q-value are provided in **Supplementary Table 22**.

## Data availability

Reference data sets used in this study are linked and annotated in **Supplementary Table 25-26**. ENCODE accession IDs for all MPRA data generated for this study are available in **Supplementary Table 26**. The UKBB analysis in this study was conducted via application number 31063.

## Code availability

Code used to analyze and visualize data related to this work will be available at https://github.com/julirsch/finemapped_mpra/.

## Acknowledgements

We thank The Jackson Laboratory Genome Technologies Core for experimental support; Jeff Vierstra for sharing unpublished data and results; Michael Stitzel, John Ray, Juan Fuxman Bass, and Stephen Rong for suggestions and conversations about the manuscript. This work was directly supported by Howard Hughes Medical Institute and by US National Institutes of Health grants UM1HG009435, R00HG010669, R00HG010669, R01HG012872, and R35HG011329. Individual support was provided by the Novo Nordisk Foundation (NNF21SA0072102) to RC and TRJ and the US National Institutes of Health grants T32GM007753 and T32GM144273 to LS. The content is solely the responsibility of the authors and does not necessarily represent the official views of the National Institutes of Health.

## Author Contributions

JCU and RT conceived the study. LS, RIC, HKF, SKR, JCU, and RT designed experiments. SK, DB and KM performed MPRA experiments. LS, RIC, HD, MK, QW, CMV, JCU, and RT performed data analysis. LS, RIC, HD, TTLN, MK, QSW, ZRM, SJC, ESL, HKF, SKR, JCU and RT interpreted results. LS, RIC, ESL, HKF, SKR, JCU, and RT drafted the manuscript. PCS, SKR, and RT secured funding, ESL, PCS, HKF, SKR, JCU, and RT supervised the study. All authors revised the manuscript and approved its final version.

