## Supplementary Figures 1-9 for "Functional dissection of complex and molecular trait variants at single nucleotide resolution"

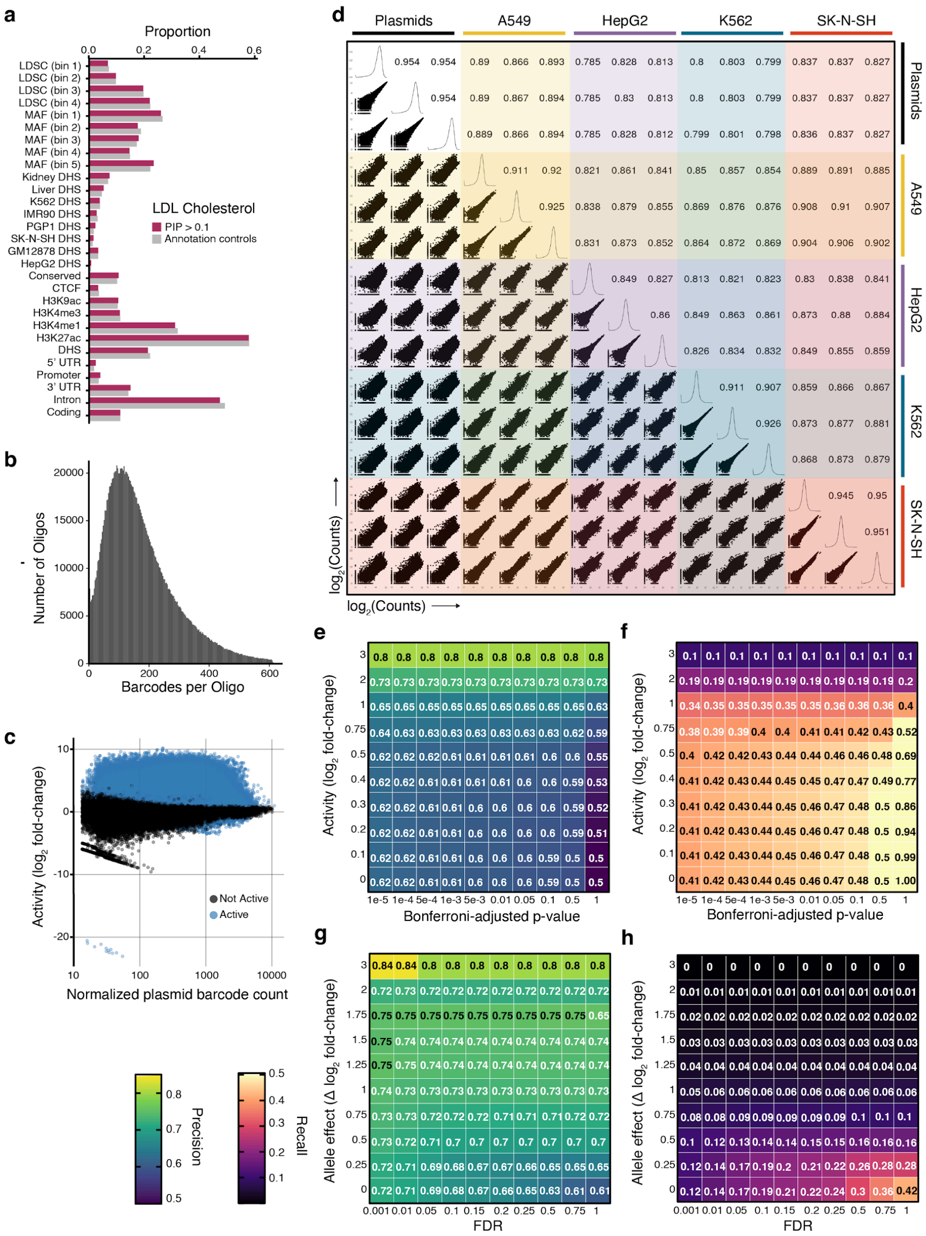


**Supplementary Fig. 1. Additional diagnostic of MPRA data quality**. **a.** Example annotation-matched control (grey) and test (PIP > 0.1) (burgundy) variant proportions in the MPRA library for LDL-C, as an exemplary trait. **b.** Distribution of barcodes per oligo across tested oligo elements. **c.** MPRA activity ($log_2$ expression RNA fold change over DNA) compared to normalized plasmid (DNA) counts. Significant elements are shown in blue (Wald's test, Bonferroni-adjusted P < 0.01) for expression differences between RNA and DNA after correcting for multiple hypothesis testing, within each cell-type/library. **d.** Correlations between normalized DNA and RNA counts ($log_2$ counts per million) across plasmids and each cell-type tested. Replicates 1, 2, and 3 (out of 5 total replicates) are shown due to space limitations. **e,f.** Precision and recall values for grid-searching across activity magnitudes and Bonferroni-adjusted p-values to determine activity thresholds. **g,h.** Precision and recall values for grid-searches across allelic effect magnitudes and FDR (at activity thresholds of magnitude > 1, Bonferroni-adjusted P < 0.01)

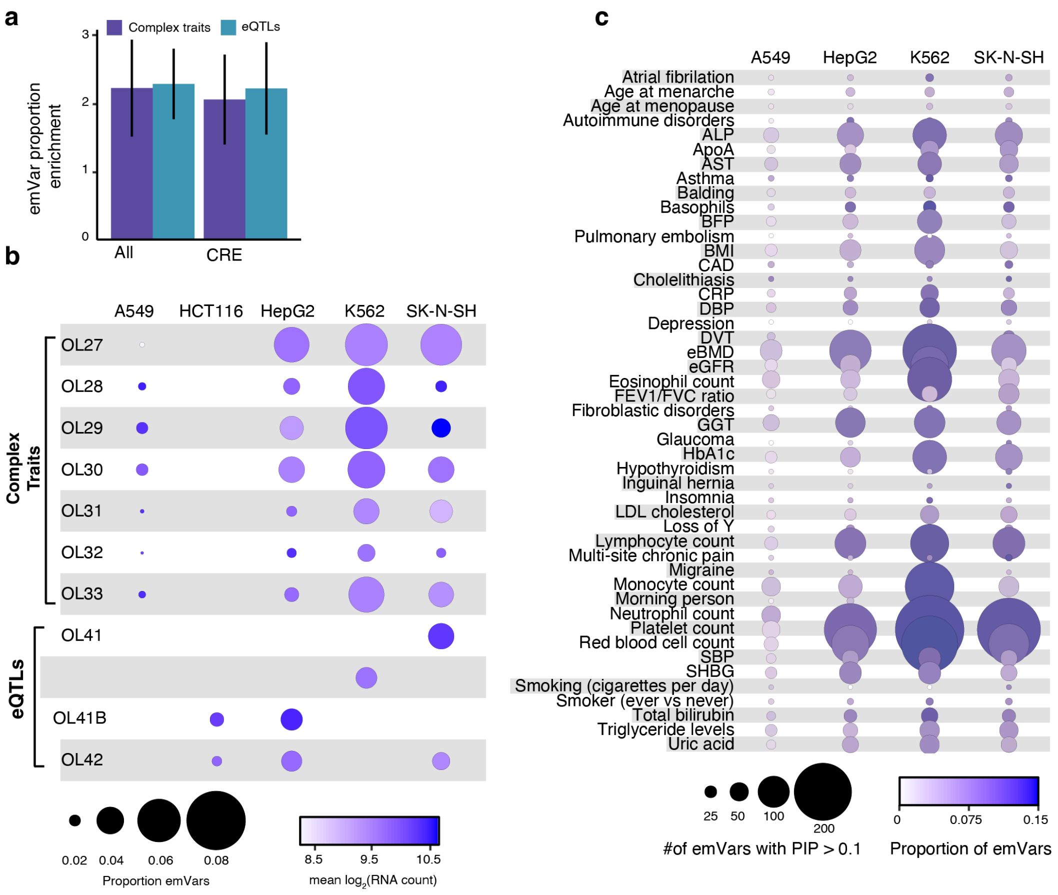


**Supplementary Fig. 2. Comparison of emVars found across libraries and cell-types. a.** Enrichment of proportions of complex trait (purple) and eQTL (blue) variants that are emVars in the highest PIP bin versus the lowest PIP bin for all variants and variants in CRE. Error bars represent 95% CIs. **b.** Proportion of variants that are emVars across cell-types and libraries. Circle size corresponds to the proportion of emVars and color indicates average RNA counts. **c.** Total number of emVars with a PIP > 0.1 (size of circle) and proportion of tested variants that are emVars (color scale) across complex traits.

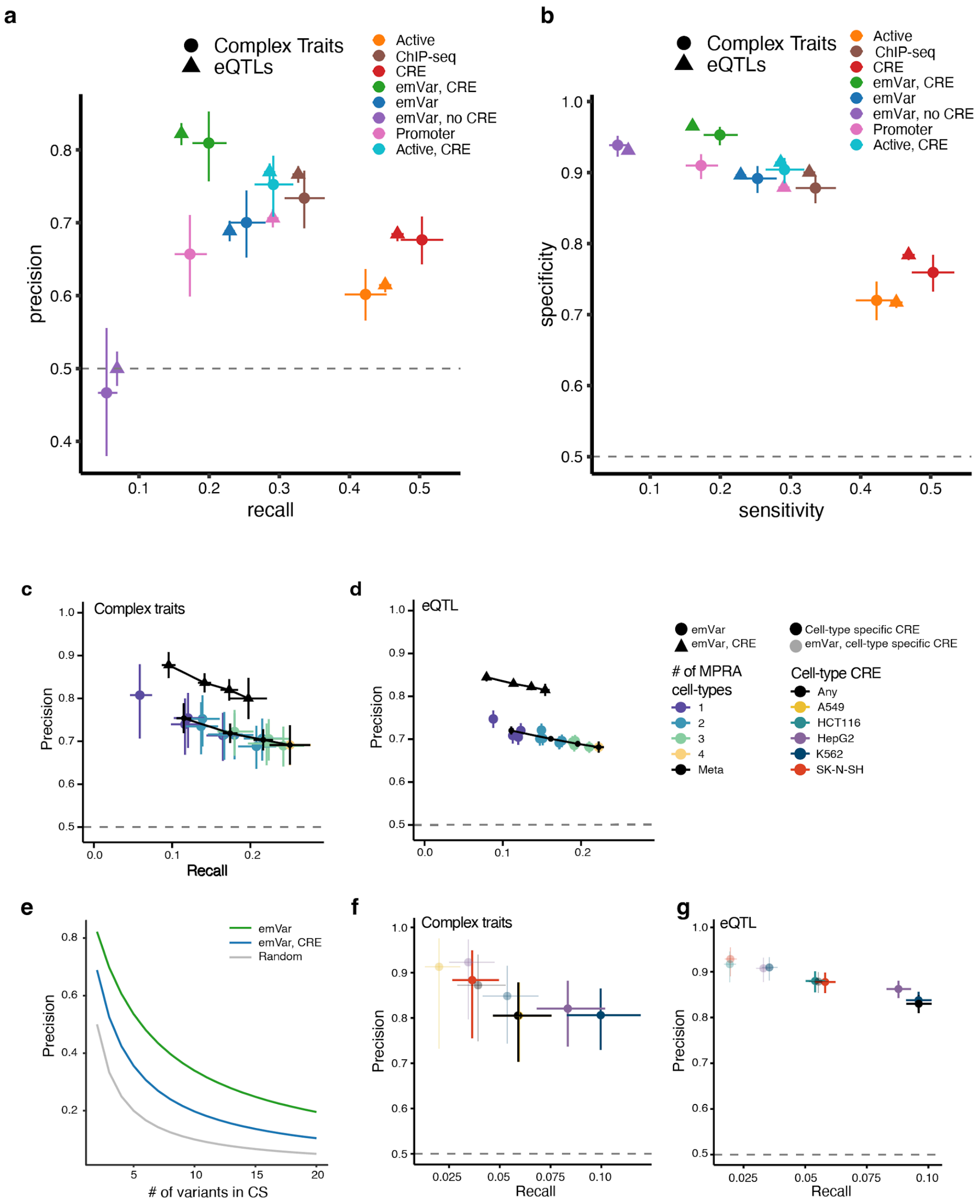


**Supplementary Fig. 3. Additional precision and recall analysis. a.** Precision-recall plots evaluating different methods for discriminating between high-confidence positive (PIP > 0.9) and negative (PIP < 0.01) variants. **b.** Sensitivity-specificity plots evaluating different methods for discriminating between high-confidence positive (PIP > 0.9) and negative (PIP < 0.01) variants. **c,d.** Precision-recall plots evaluating different methods (emVar, emVar andCRE intersection) for discriminating between positives (PIP > 0.9) and negative (PIP < 0.01) and complex trait (**c.**) and eQTL (**d.**) variants, varying the number of cell-types included. Meta-analysis points across each number of included cell-types are shown in black. **e.** Precision of each annotation across CS size The background rate (1 / [CS size - 1]) is shown in grey and compared to emVars alone (green) or emVars in CRE annotations (blue). **f,g.** Precision-recall plots evaluating the utility of cell-type specific CRE annotations on their own versus the intersection of cell-type specific CREs and emVar measurements in discriminating between high-PIP (PIP > 0.9) and control (PIP < 0.01) complex trait (**f.**) andeQTL (**g.**) variants. Meta-analysis points are indicated in black.

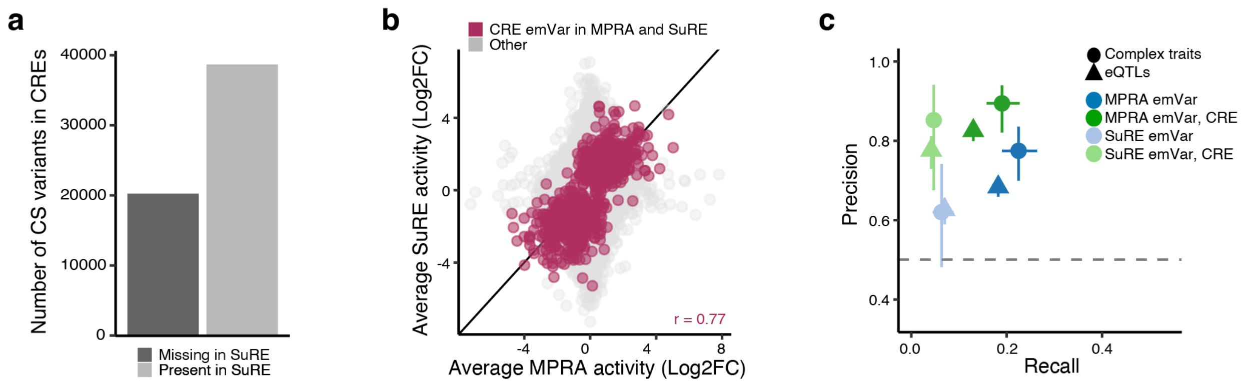


**Supplementary Fig. 4. Comparisons to previous reporter assays. a.** Number of CS variants in CREs for the traits investigated in this study that were tested previously in SuRE (light grey) or are missing (dark grey). **b.** Pearson correlation between variant activity ($\log_2$[fold-change])estimates in SuRE compared to MPRA. Variants in CREs that are significant emVars in both studies at the same reported FDR level (< 0.05) are labeled in burgundy. **c.** Precision-recall plots evaluating MPRA emVars or SuRE emVars, with or without a CRE filter for discriminating between high-confidence positive (PIP > 0.9) and negative (PIP < 0.01) variants.

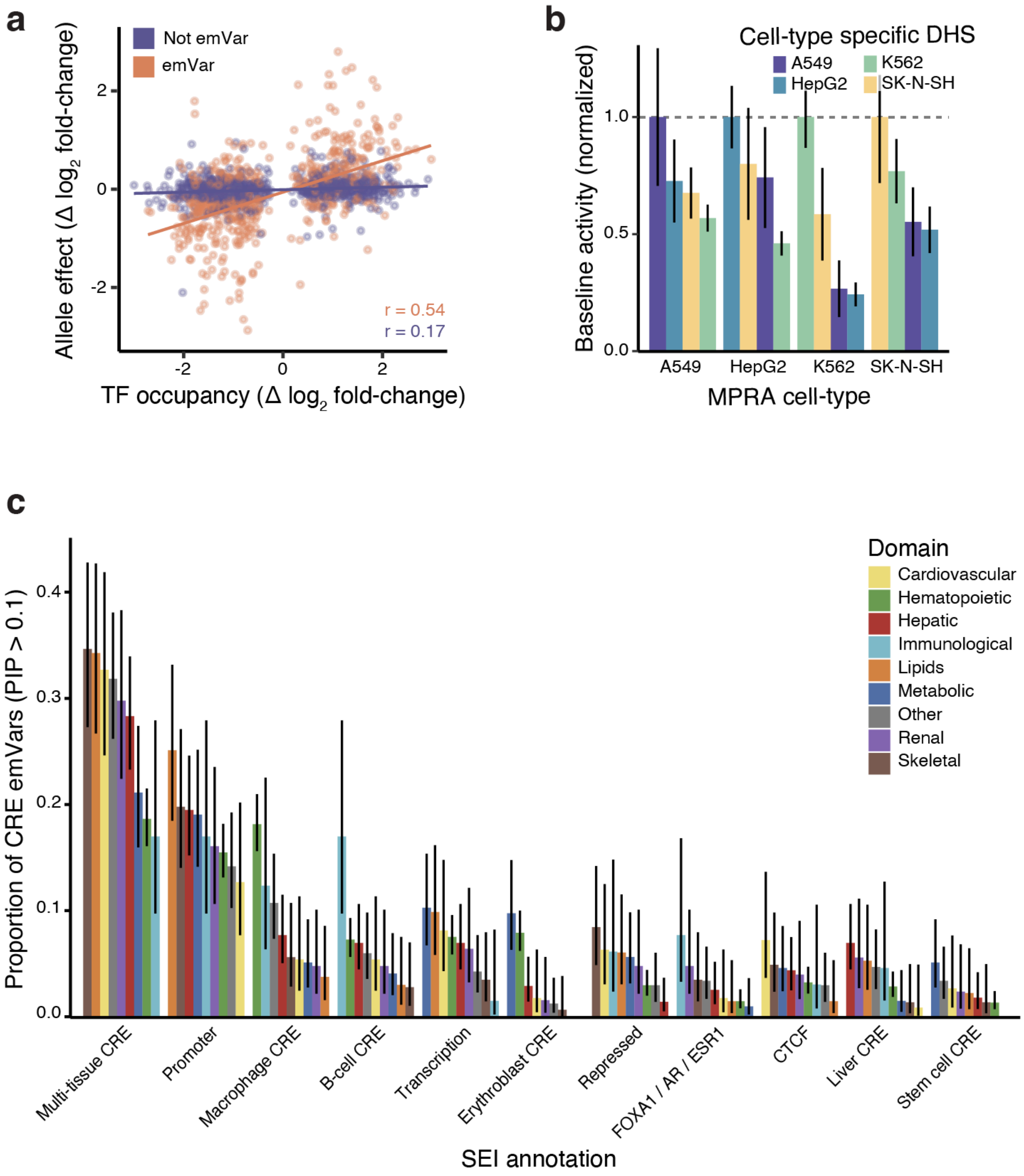


**Supplementary Fig. 5. Additional evidence of reporter assays recapitulating regulatory function. a.** Transcription factor occupancy QTL effects sizes are correlated (Pearson r) with MPRA allelic effects. emVars are shown in orange and non-emVars in purple. **b.** Baseline activity is highest for variants in cell-type specific DHS that match the cell-type assayed. Baseline activity is normalized to 1 for the reference cell-type. **c.** Complex trait variants falling in each of the SEI categories were stratified into domains. Across each domain, the proportion of CRE emVars falling into each SEI category was calculated. Error bars represent 95% CIs.

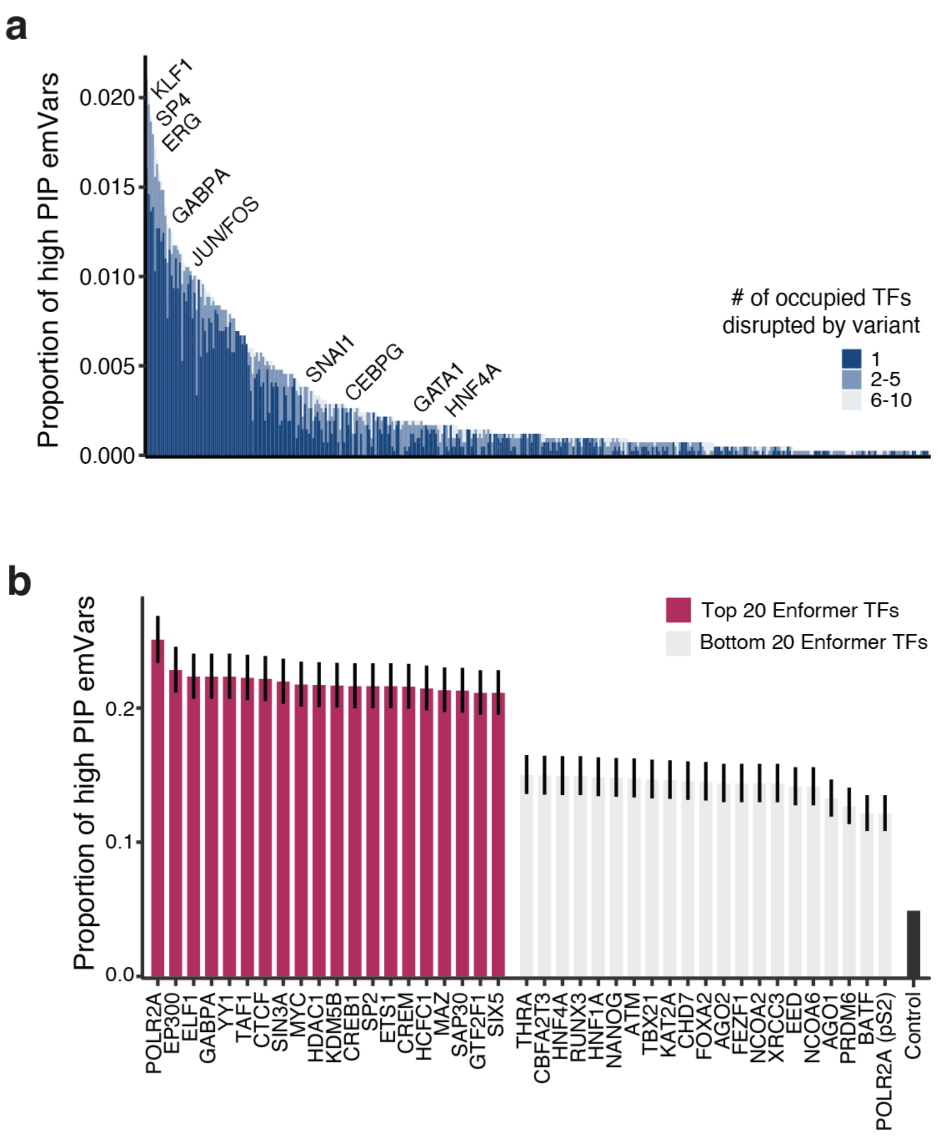


**Supplementary Fig. 6. TF disruptions underlying emVars. a.** Proportion of likely causal variants that disrupt an occupied transcription factor binding motif. Variants often disrupt multiple occupied motifs, which is indicated by color. **b.** Proportion of high PIP CRE emVars that disrupt various TF occupancies as predicted by Enformer. The top and bottom 20 TFs enriched at high PIP emVars are shown alongside the low PIP, non-emVar control. Error bars represent 95% CIs.

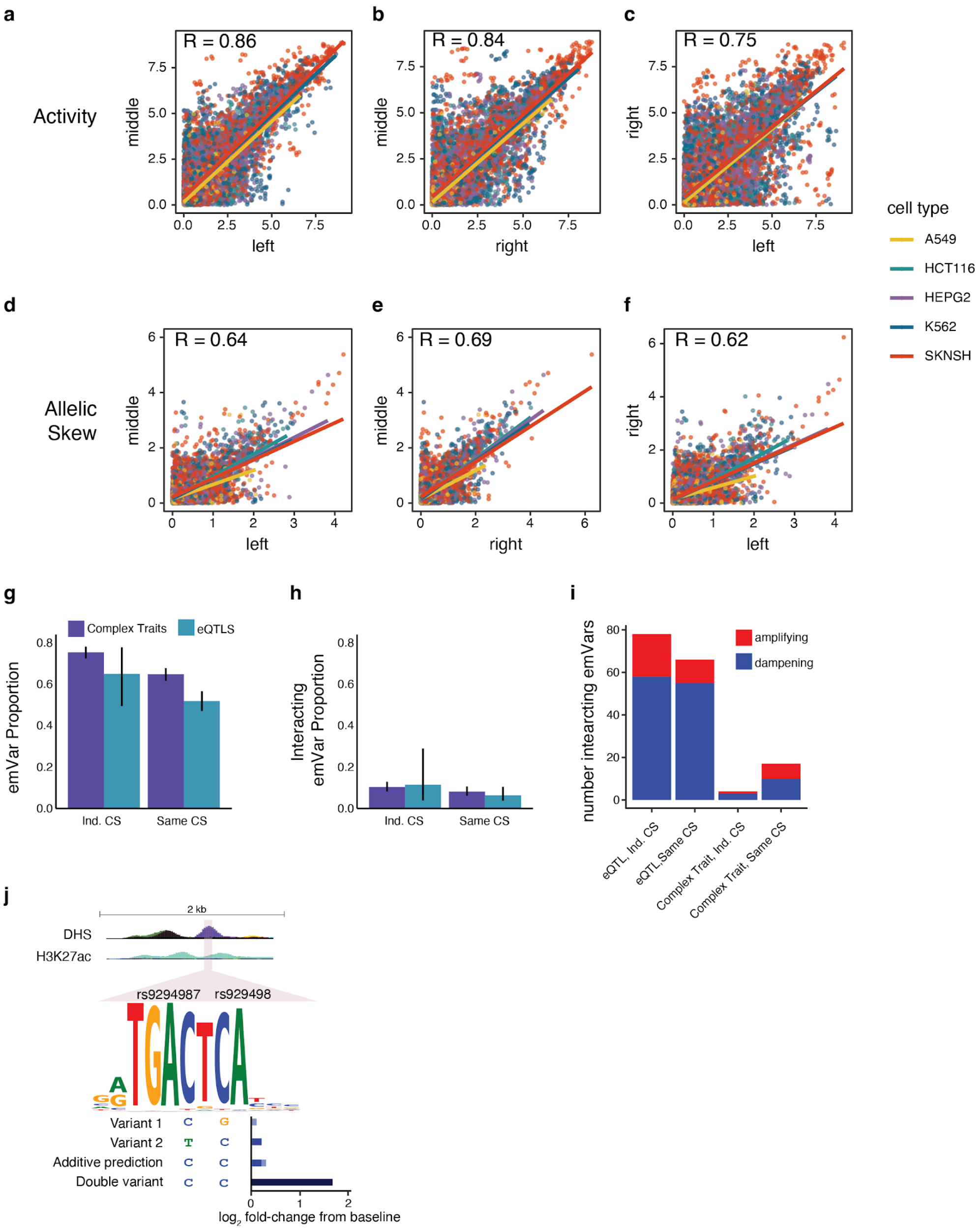


**Supplementary Fig. 7. Additional evidence for regulatory allelic heterogeneity and multiple causal variants. a-c.** Scatter plots of activity from all 4 diplotypes across 3 windows. Each point represents an activity measurement for one diplotype in one cell-type. Pearson correlations are shown for all points indicated in the plot. Lines represent linear fits of activity between windows stratified by cell-type. **d-f.** Same as **a-c.**, except for all 3 allelic effects (compared to the homozygous reference). **g.** Proportion of emVars across any diplotype stratified by independent vs same CS and complex trait vs eQTL variants. Error bars represent 95% CIs. **h.** Proportion of non-additive emVars across any diplotype stratified by independent vs same CS and complex trait vs eQTL variants. Error bars represent 95% CIs. **i.** Distribution of amplifying versus dampening interacting pairs across eQTLs and complex traits stratified by independent vs same CS and complex trait vs eQTL variants. **j.** Example of an amplifying non-additive variant pair, rs9294987 and rs9294988 (shaded purple), which are associated with systolic blood pressure. The closest gene is *THBS2* and the variant pair jointly improve a Jun binding motif. Additive prediction is the sum of the re-coded allelic effects from variant 1 (TG vs CG) and variant 2 (TG vs TC). Double variant is the observed difference between TG and CC. (FDR = $2.0 \times 10^{-40}$).

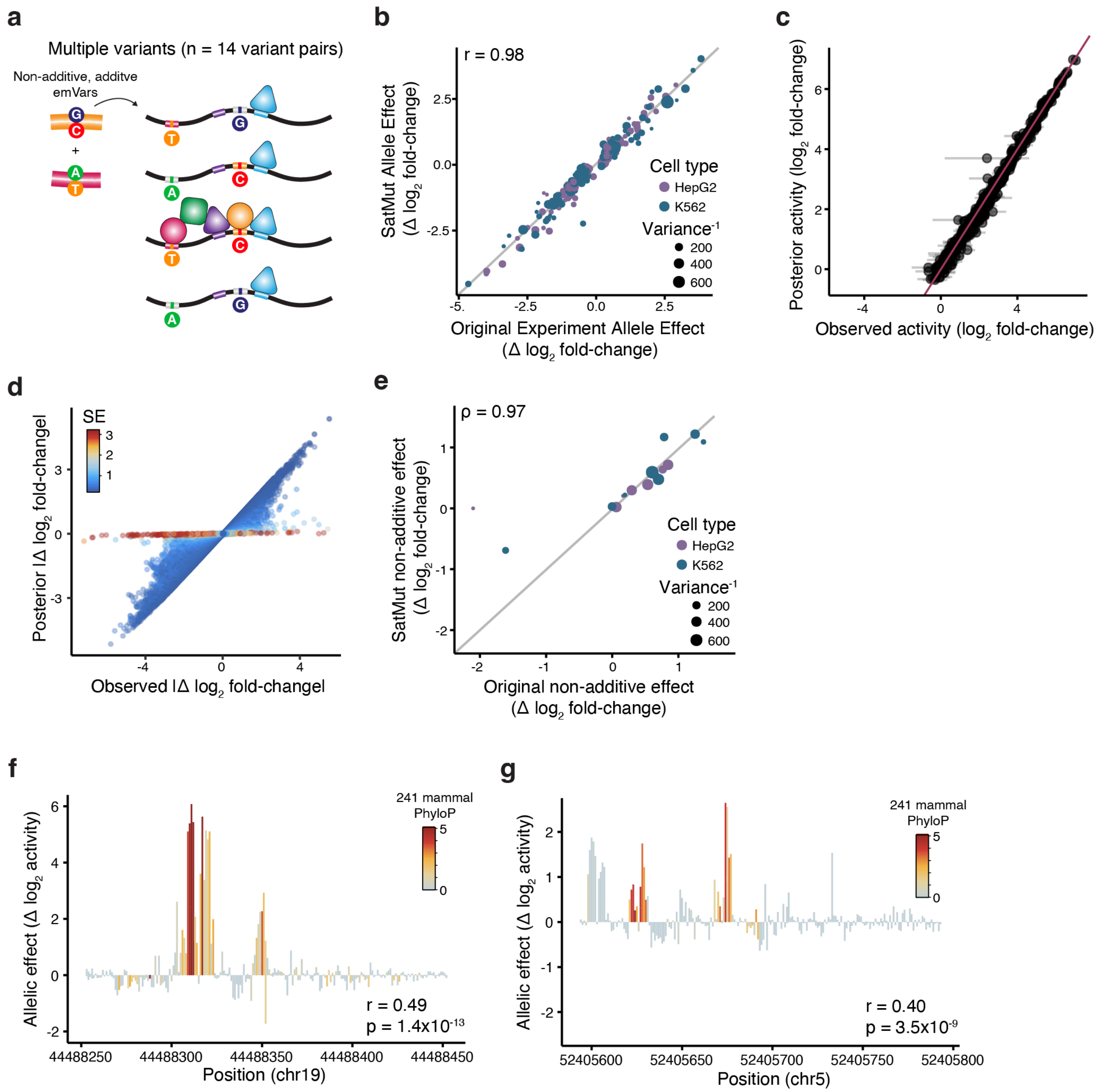


**Supplementary Fig. 8. Saturation mutagenesis recapitulates allelic effects. a.** Schema for 14 variant pairs tested with SatMut across all four possible diplotypes. **b.** Scatter plot between emVar allelic effects measured in the original experiment and in SatMut. Measurements are shown for HepG2 (purple) and K562 (blue) with circle size indicating inverse variance (IV). The Pearson correlation coefficient is shown. **c.** Comparison of raw SatMut baseline activity and empirical Bayes-adjusted baseline activity for all 284 background sequences. Error bars represent 95% CIs. **d.** Comparison of raw SatMut baseline activity and empirical Bayes-adjusted allelic activity across all substitutions. Substitutions with high SEs have allelic effects shrunk towards zero. **e.** Scatter plot between allelic pairs' interaction effects measured in the original experiment and in SatMut, plotted identically to **b**. The Spearman correlation coefficient is shown instead of Pearson given the small sample size. **f.**,**g.** Correlation between the average allelic effect (contribution) and mammalian constraint (PhyloP) for two separate loci. Positive values indicate activating sequences whereas negative values indicate repressive sequences.

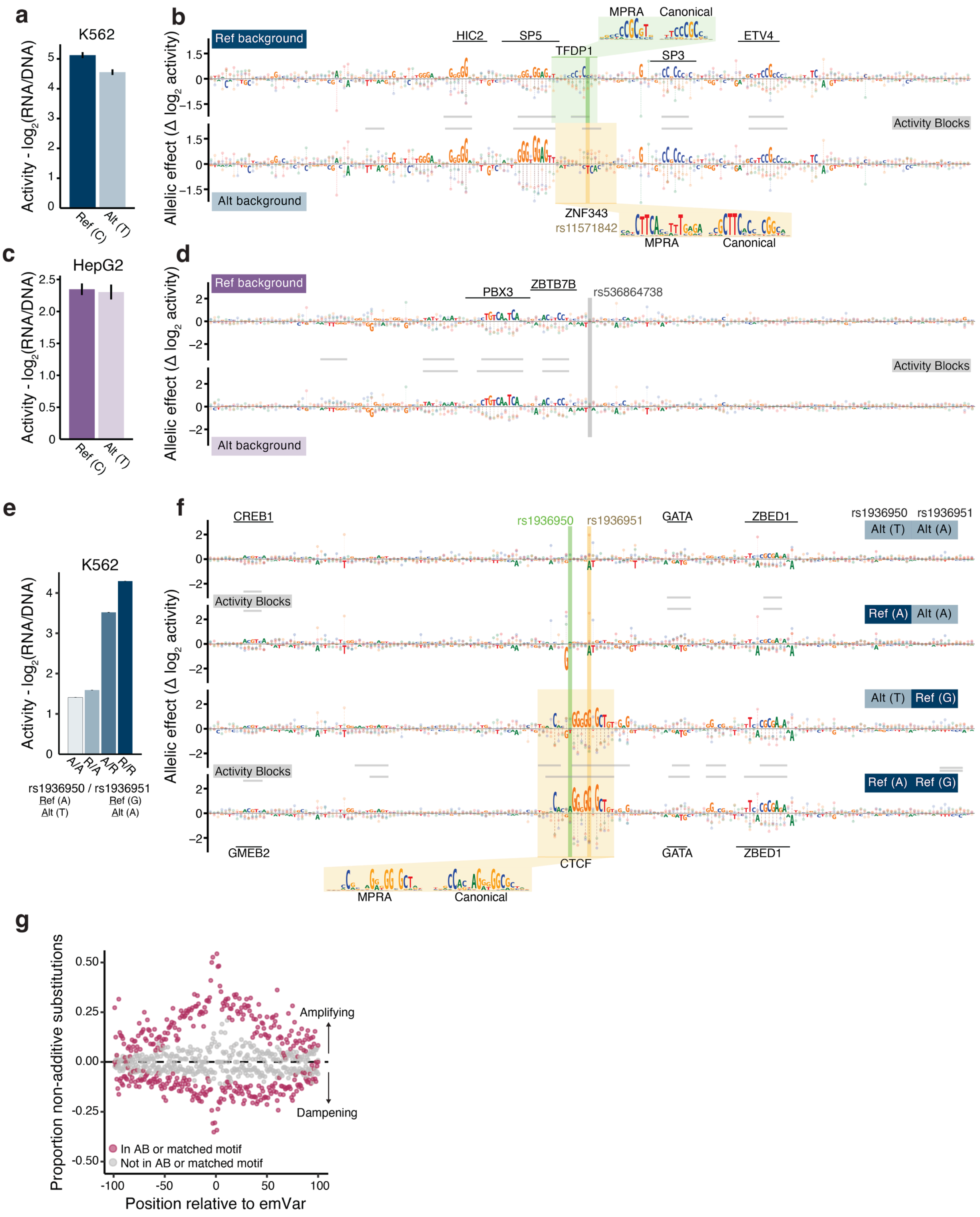


**Supplementary Fig. 9. Additional saturation mutagenesis examples. a.,c.,e.** MPRA results of transcriptional activity for the reference (darker shade) and alternative (lighter shade) allele(s) in either K562 (blues) or HepG2 (purples) for emVars rs11571842 (**a.**), rs536864739 (**c.**), and rs1936950/rs1936951 (**e.**). Error bars indicate SEs. **b.,d.,f.** Nucleotide contribution scores across the 200 bp elements containing emVars from **a.**, **c.**, and **e.** are highlighted by a dark yellow or green bar. Activity measurements for all positions when tested on the reference (*top*) or alternative (*bottom*) allele for **b.**,**d.** or haplotype combinations in order of activity levels (**f.**) are depicted as lollipops indicating the change from baseline activity (Δ $\log_2$ activity). Activity blocks (ABs) are labeled with a gray bar and matching TF motifs are highlighted with a black bar. Shaded boxes overlap allele(s) of interest, with a callout of the SatMut constructed motif (MPRA) and canonical motif PWM (Canonical). **g.** Scatter plot of variant pair interactions between the centered emVar and all other substitutions made by SatMut. Plotted is the proportion of non-additive substitutions (y-axis) that are amplifying (positive) or dampening (negative) with respect to the distance of the substitution to the emVar (x-axis). Interactions are separated by whether the adjacent substitution resides in an activity block (AB) or SatMut matched motif.

### Supplementary Tables

1. Summary of Variant Fine-Mapping
2. Variant Oligo Sequences
3. MPRA Results
4. Test and control variants for precision and recall plots
5. Precision and Recall Results
6. Precision simulations across varying CS sizes
7. High Confidence Variants: emVars in CREs in non-coding CSs
8. High Confidence emVars associated with an additional 512 Human Diseases
9. Transcription Factor enrichments at high-PIP emVars vs emVars
10. Cell Type Specific Activity and Allelic Skew from mashr
11. SEI enrichment and proportion analysis
12. Occupied TF Motif Disruptions by High-PIP emVars
13. Correlations between Allelic Skew and TF Motif Disruption
14. Oligo sequences for variant pairs across diplotypes and windows
15. Credible Set status and annotation for variant pairs
16. Overview of Variant Pair MPRA Results
17. Summary of non-additive (interacting) variant pairs
18. Multiple Causal Variant results across CS sizes and LD thresholds
19. Summary of Saturation Mutagenesis Motif and TF Occupancy Annotations
20. Summary of Saturation Mutagenesis Results
21. Saturation Mutagenesis enrichment of motifs at activity blocks (ABs)
22. Correlations between Saturation Mutagenesis MPRA results and PhyloP scores
23. Oligo Sequences of MPRA Controls
24. Primer sequences
25. Columns included in CRE definition and CRE binary
26. Data sources for data generated or used in this manuscript
